# When good species have porous boundaries: weak reproductive isolation and extensive gene flow between *Mimulus glaucescens* and *M. guttatus* in northern California

**DOI:** 10.1101/2022.09.08.507212

**Authors:** C. T. Ivey, N. M. Habecker, J. P. Bergmann, J. Ewald, J. M. Coughlan

## Abstract

Barriers to reproduction are often how progress in speciation is measured. Nonetheless, a key unresolved question is the extent to which reproductive barriers diminish gene flow in incipient species in nature. The Sierra Nevada foothill endemic *Mimulus glaucescens* and the widespread *M. guttatus* are considered to be distinct species based on contrasting vegetative traits, but barriers to reproduction are not readily apparent, although these species are not known to hybridize in nature. To explore boundaries between taxa, we examined 15 potential reproductive barriers between species in a Northern California area of broad sympatry. Most barriers, with the exception of ecogeographic isolation, were weak, and total isolation for each species was estimated to be incomplete. Population genomic analyses of range-wide and broadly sympatric accessions revealed that gene flow between these taxa is common across the range, and rampant within areas of sympatry. Thus, despite fairly strong ecological differentiation - which may be involved in maintenance of vegetative differences - ecological isolation is a weak barrier to gene flow in this system. This work underscores the value of combining classical measures of reproductive isolation with estimates of natural gene flow for studies of speciation in natural communities.

## INTRODUCTION

Species boundaries are often considered to be maintained by reproductive barriers that limit interbreeding (Mayr 1942). The effectiveness of these barriers at limiting gene flow typically varies (Lowry et al. 2008a; Widmer et al. 2009; Baack et al. 2015), and in many cases boundaries are established only through multiple barriers in combination, no one of which is sufficient to preclude all gene flow (Ramsey et al. 2003; Kay 2006; Ostevik et al. 2016; Christie and Strauss 2019). Species boundaries also can be maintained with few barriers to reproduction if opposing selection is sufficiently strong (Via 2009; DiVittorio et al. 2020). However, interbreeding can itself alter the strength of reproductive barriers, either positively or negatively (Servedio and Kirkpatrick 1997; Runemark et al. 2019). Moreover, the pace and sequence of barrier evolution appears to be quite variable (Widmer et al. 2009; Baack et al. 2015; Coughlan and Matute 2020), and in some cases results in apparently stable partial isolation, in which some gene flow is routine (Stankowski and Ravinet 2021).

As a consequence, some authors have suggested that boundaries between species or populations are best regarded as falling along a continuum, with panmixia and complete isolation at the extremes, and the strength of reproductive barriers between two populations defining their position on the continuum (Merill et al. 2011; Powell et al. 2013; Feder et al. 2014; Harrison and Larson 2014; Christie and Strauss 2019; Stankowski and Ravinet 2021). In this context, estimates of reproductive isolation between species or populations might be expected to inform predictions about genetic divergence as well as the likelihood of ongoing gene exchange. Estimates of barrier strength, however, are not always reliable indicators of gene flow. Sambatti et al. (2012), for example, estimated reproductive isolation between *Helianthus annuus* and *H. petiolaris* to exceed 0.999, based on multiple potential barriers, but nonetheless found extensive introgression between species, based on sampling multiple nuclear loci. De La Torre et al. (2014) similarly found extensive admixture between two *Picea* species based on genome-wide SNP panels, but also found evidence that adaptation to different climatic conditions helped maintain species integrity. Finally, although prezygotic reproductive isolation has been estimated to exceed 0.99 between the recently diverged *Mimulus guttatus* and *M. nasutus* (Martin and Willis 2006), introgression appears common between these taxa (Brandvain et al. 2014; Kenney and Sweigart 2016). Thus, interpretation of estimates of reproductive isolation with respect to species boundaries benefits from more direct estimates of gene flow and introgression based on genomic sampling.

Progress in understanding the evolution of reproductive isolation and patterns of speciation has often come from studies of groups of related taxa (Coyne and Orr 1989; Moyle et al. 2004; Owens and Rieseberg 2013; Christie and Strauss 2018; Coughlan and Matute 2020). One such group includes the diverse wildflower genus *Mimulus* (Wu et al. 2008; Twyford et al. 2015; Nelson et al. 2021; note that some taxa were placed into other genera by Barker et al. [2012], but we herein follow Lowry et al. [2019] in the use of *Mimulus* given the lack of a well-resolved phylogeny). Comparative studies involving *Mimulus* have suggested habitat divergence, mating system differentiation, or shifts in floral traits associated with pollination may be common during speciation (Grossenbacher and Whittall 2011; Grossenbacher et al. 2014; Sobel et al. 2014). As divergence proceeds, however, post-pollination barriers appear to increase in strength (Ferris et al. 2014; Coughlan et al. 2020; Sandstedt et al. 2021; Joffard et al. 2021). Introgression has also been a salient feature during speciation in *Mimulus* (Brandvain et al. 2014; Kenny and Sweigart 2016; Nelson et al. 2021). Within the adaptive radiation of *Mimulus* section *Erythranthe*, for example, extensive ancestral and recent introgression has been documented, even between taxa with strong reproductive barriers, which underscores the importance of natural selection in boundary maintenance (Nelson et al. 2021).

Many of the *Mimulus* taxa studied to date have relatively conspicuous contrasts in distribution, habitat, floral traits, or mating system that result in barriers to reproduction. In contrast, the taxa studied herein, *M. glaucescens* and *M. guttatus*, have few apparent constraints on gene exchange (see below). Taxonomists, however, have treated them as distinct species based on marked contrasts in vegetative morphology (Fig. 1; Baldwin et al. 2012; Barker et al. 2012; Nesom 2012), and natural hybrids are not commonly observed (Vickery 1964). Here, we characterize 15 potential reproductive barriers and use whole-genome resequencing to estimate barrier strength and compare patterns of reproductive isolation with patterns of population genetic structure, divergence, and introgression to better understand boundaries between these taxa. Our results contribute to understanding how reproductive barriers relate to introgression and illuminate the speciation process in recently diverged taxa.

**Figure 1:**
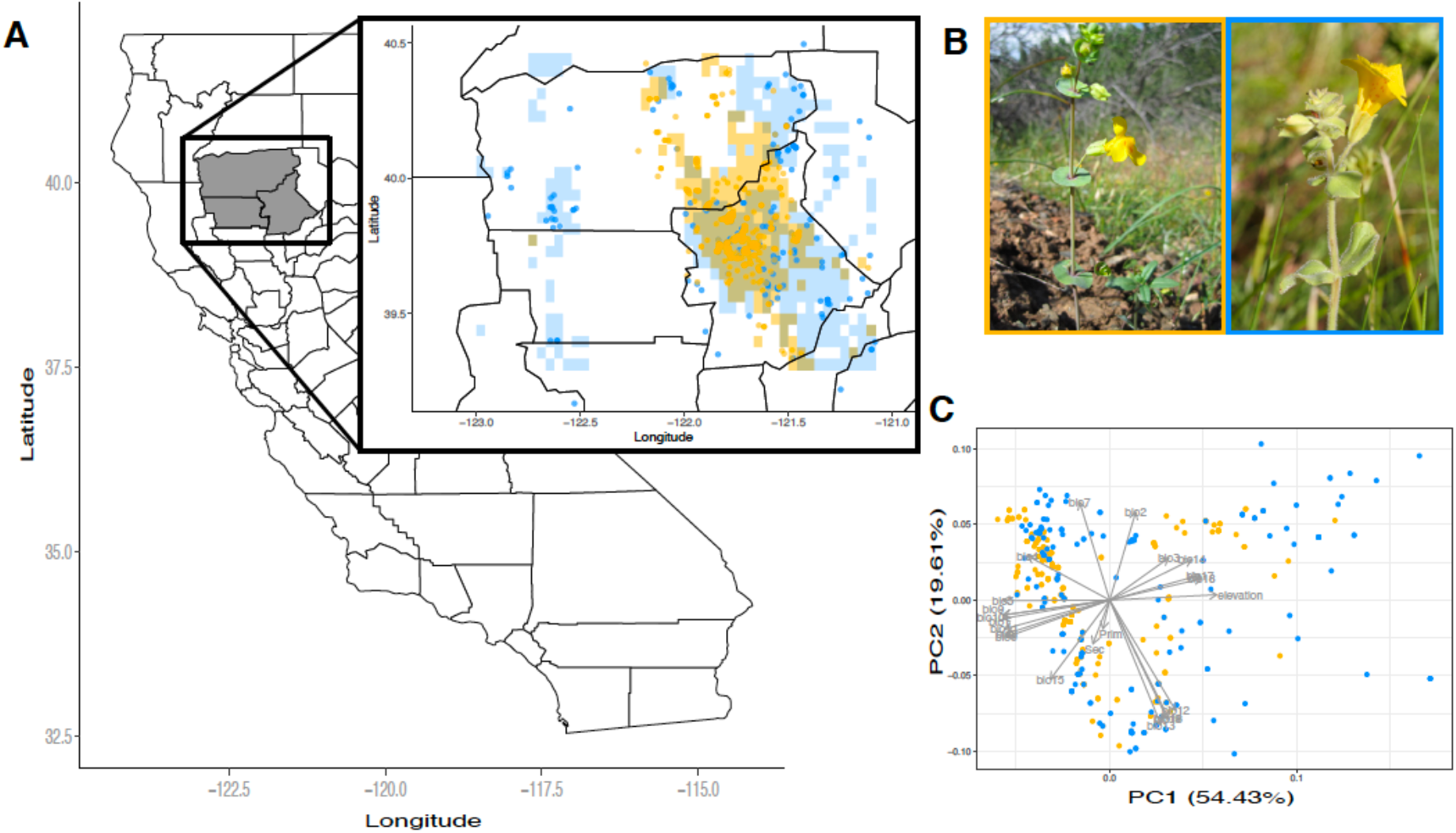
Geographic distributions of *Mimulus glaucescens* and *M. guttatus*. (A) State of California with focal counties in gray; inset: predicted local ranges of *M. guttatus* (blue) and *M. glaucescens* (yellow) based on maxent modeling. Points represent collection records upon which the models are based. (B) *M. glaucescens* (left) and *M. guttatus* (right), showing distinct vegetative morphology, (C) PCA based on 19 Bioclim variables, elevation, and two soil substrate variables used to build the maxent models.

## METHODS

### Study system

The genus *Mimulus* (family Phrymaceae) is a diverse group of at least 180 plant species (Grant 1924; Beardsley et al. 2004; Barker et al. 2012) that has been a focus for studies of evolution and speciation (Schemske and Bradshaw 1999; Martin and Willis 2007; Lowry et al. 2008b; Wu et al. 2008; Sobel 2014; Twyford et al. 2015; Nelson et al. 2021; Sandstedt et al. 2021). Some taxa within section *Simiolus*, which includes *M. guttatus*, have been taxonomically problematic (Munz 1959; Hickman 1993; Nesom 2012) and likely represent multiple recently evolved lineages (Beardsley et al. 2004; Sweigart and Willis 2003; Lowry et al. 2008b; Barker et al. 2012). The similar *M. glaucescens*, however, has been considered distinct from *M. guttatus* based on vegetative characters (Baldwin et al. 2012; Nesom 2012). Nesom (2012) described *M. glaucescens* as “clearly defined” in a recent treatment of the section. *Mimulus glaucescens* has peduncular bracts that are perfoliate to connate, glabrous, distinctly glaucous, and with entire margins, whereas the peduncular bracts of *M. guttatus* are generally distinct, bearing trichomes that are often glandular, lacking in glaucousness, and with margins that are often dentate (Fig. 1; Baldwin et al. 2012). Phylogenies suggest the two are closely related, and possibly sister taxa (Beardsley et al. 2004; Lowry et al. 2019). Both species are mixed-mating herbs, flower during the spring, occur in similar habitats, and are *n* = 14 (Vickery 1978; Ritland and Ritland 1989; Baldwin et al. 2012). *Mimulus glaucescens* is endemic to a few counties within northern California, whereas the broad western North American range of *M. guttatus* wholly overlaps that of *M. glaucescens* (Baldwin et al. 2012; Nesom 2012). Within the range of *M. glaucescens, M. guttatus* is found on non-serpentine soils (C. T. Ivey, pers. obs.), although it occurs on serpentine elsewhere (Selby and Willis 2018; Sianta & Kay 2021). In contrast, *M. glaucescens* occurs on both serpentine and non-serpentine soils (Nesom 2012). Populations in the current study were from non-serpentine sites unless otherwise noted. Previous work using few maternal lines reported reduced hybrid seed set and F_1_ fertility between *M. glaucescens* and *M. guttatus* (Vickery 1964). Otherwise, little appears to be known about what limits hybridization between these taxa.

### Pre-pollination barriers

To characterize the importance of pre-pollination barriers to species boundaries, geographical distribution, local habitat, flowering phenology, floral visitor behavior, and floral morphology were compared between species. Habitat associated with each species was modelled using 2512 georeferenced collection records from the Global Biodiversity Information Facility, which were filtered after download (GBIF; 2194 *M. guttatus* records: https://doi.org/10.15468/dl.ywy9s3 and 318 *M. glaucescens:* https://doi.org/10.15468/dl.r93avz; see Supplemental Methods and Results) and environmental data including primary and secondary geological substrates, elevation, and 19 bioclimatic variables (https://www.worldclim.org/), all standardized to a resolution of 2.5 minutes. Species distribution models were constructed using MaxEnt by sampling locations of collection records as well as 1000 randomly located background samples from across the geographic extent to create pseudo-absence data (Coughlan et al. 2021). Models were computed separately for each species, and reproductive isolation was inferred based on the number of shared and unshared pixels in the map following equation RI_4C_ from Sobel and Chen (2014) (see Supplemental Methods and Results).

In addition, local-scale habitat characteristics were measured for each species within a 6 km^2^ region of sympatry along Butte Creek within Butte county California. Within each of 15 populations (*M. glaucescens* = 9; *M. guttatus* = 6), slope, percent shade cover, percent vegetative cover, percent bare ground, percent debris, and percent standing water was measured for each of 8-10 randomly chosen plants (*sensu* Li et al. 2018). Scores of the first two principal components of the combined data were compared between species using a mixed-model ANOVA with species as the fixed effect and population as the random effect.

Flowering phenology was compared between species within a flowering season for four populations of each species occurring within a 2 km^2^ region along Butte Creek (Table S1). The number of flowering plants was recorded once each week from mid-March (when no plants were flowering) until early August (after all flowering had stopped). Flowering plant numbers were compared between species using a repeated measures model in which plant species, day of observation, and their interaction were fixed effects, and population pair was a random effect. The day of observation was the repeated effect with population as the subject.

The behavior of floral visitors was compared between species using experimental mixed-species arrays placed near natural populations of each species (Table S1). Each array consisted of 18 plants of each species (36 total) in 9 cm^2^ pots arranged into a 6 × 6 grid at 0.5 m spacing. Each plant in the array had a single first-day flower open during observations. Arrays were observed between 0900 and 1500 for 1 hr intervals (total 67 hr) during which the visitor taxon and sequence of visited plants was recorded. The likelihood of floral visitors moving conspecifically (response = 0) vs. heterospecifically (1) was compared for visitors foraging on each species using a generalized linear mixed model with a binomial error distribution. Fixed effects included the plant species from which the visitor moved, site location, and their interaction. Random effects included visitor taxon (Table S2) and the observation.

First-day floral morphology was compared between species using greenhouse-grown plants reared from open-pollinated seeds collected in four populations of each species (Table S1). Corolla width at the widest point, corolla tube width at distal opening, anther length, pistil length, and herkogamy were measured to the closest mm for 120 first-day flowers (*M. glaucescens n* = 73; *M. guttatus n* = 47), and scores of the first two principal components based on family mean values were compared between species using a mixed-model ANOVA with species as the fixed effect and population as the random effect (see Supplemental Methods and Results).

### Post-pollination barriers

To characterize the importance of post-pollination barriers to species boundaries, pollen-pistil interactions, seed number, seed germination, and several measures of post-germination performance were measured following controlled hand-pollinations of emasculated flowers within species (both within- and between-populations) and between species. Hand-pollinations were performed on plants reared in the greenhouse from open-pollinated seeds collected from at least two populations of each species within the Butte Creek watershed (Table S1).

To evaluate pollen-pistil interactions, pollen adhesion, pollen germination, and pollen-tube growth rate (PTGR) were compared among types of hand-pollinations (*sensu* Kay 2006). The success of pollen adhesion was compared using a modification of Zinkl et al. (1999). A known number of pollen grains was applied to first-day stigmas (*M. glaucescens n* = 42; *M. guttatus n* = 53), which were then vortexed in a phosphate-Tween-20 solution, centrifuged for 60 sec at 7200 rpm, and fixed in 9:1 ethanol:glacial acetic acid for 24 hr before staining with basic fuchsin and viewing at 100x to record number of adhered pollen grains. Pollen germination was observed in styles (*M. glaucescens n* = 61; *M. guttatus n* = 56) that were excised 6-12 hr following hand-pollination, fixed and stained in decolorized 0.1% aniline blue before viewing at 100x under epifluorescence. The number of pollen tubes in styles and number of pollen grains on stigmas was recorded. Pollen tube growth rate was observed in styles (*M. glaucescens n* = 25; *M. guttatus n* = 42) that were excised five minutes after hand-pollination and placed onto enriched 1% agar. Style length was measured to the nearest 0.1 mm using a dissection microscope fitted with an ocular micrometer and the time in min for pollen tubes to emerge from the style was recorded (see Supplemental Methods and Results).

Seed number was recorded for individual fruits resulting from hand-pollinations of emasculated flowers (*M. glaucescens n* = 119; *M. guttatus n* = 185) using digital images following Ivey and Carr (2012). Sixty fruits from each species were randomly chosen, and 20 seeds from each were scored for germination (number of seedlings) two weeks after sowing into 100 cm^2^ pots. One randomly chosen seedling from each fruit was grown to maturity, and the number of days until the first flower opened (DFF) was scored to indicate development.

In addition, ovule number and pollen viability were scored from the first open flower following Carr and Dudash (1997). After 59 days, total number of flowers was recorded, and all above-ground biomass was dried to constant mass at 55º C before weighing to nearest mg.

Post-pollination traits were compared using mixed-model ANOVA that included hand-pollination category (within-population, between-population conspecific, and heterospecific), the species of pollen recipient, and their interaction as fixed effects, whereas random effects included the population identities of pollen donor and pollen recipient. Additional details about models used for each trait can be found in Supplemental Methods and Results.

### Reproductive isolation

Reproductive isolation was estimated for each of the potential barriers above where possible, following Sobel and Chen (2014). To provide context for interpreting estimates of isolation between species, estimates of isolation were calculated between populations within each species as well (Table S3). Total reproductive isolation was estimated from all barriers using equation RI_4E_ from Sobel and Chen (2014). Stability of the estimates was evaluated by constructing 95% confidence intervals for individual barriers as well as total isolation using 10,000 bootstrap replicates except in the case of ecogeographic isolation in which we used 1,000 bootstraps for computational efficiency (*sensu* Ostevik et al. 2016; Christie and Strauss 2019).

### Genomic analyses

We paired previously published whole-genome re-sequencing data (Brandvain et al. 2014, Puzey et al. 2018, Coughlan et al. 2020, Coughlan et al. 2021) with new whole-genome re-sequencing data described herein to assess taxonomic relationships, patterns of population genetic structure, and the presence of introgression between *M. guttatus* and *M. glaucescens* (See Table S4 for sample details). In total, we leverage data from 75 accessions representing 10 named species across the *M. guttatus* species complex plus a single accession of *M. dentilobus* as an outgroup. For the newly described whole-genome re-sequencing data, we followed protocols described elsewhere for sample preparation, sequencing, and preparation of VCF files (e.g. Coughlan et al. 2020, Coughlan et al. 2021; see Supplemental Methods for details).

#### Phylogenetic relationships

To assess phylogenetic relationships among named members of the *M. guttatus* species complex we inferred a Maximum Likelihood (ML) phylogeny as well as 1,000 ultrafast bootstraps using IQtree V2.0.3 (Nguyen et al. 2015) with a TVM+F+R4 model, which IQtree identified as the best fitting model based on both AIC and BIC. For this analysis, we filtered the VCF file to include sites in which 72 of the 75 accessions were genotyped (~95% of accessions), which resulted in 195,771 sites, 146,697 of which were parsimony informative. We also performed principal components analyses on genome-wide data using PCANGSD (Korneliussen et al. 2014) for all members of the complex (Fig. S1) as well as just *M. guttatus* and *M. glaucescens*, to evaluate whether named species correspond to genetically distinct clusters. We furthered this analysis by performing K-means clustering analysis using both PCANGSD (in which K is identified as the number of significant PCs +1) and more traditional K-means clustering analyses using NGSAdmix (Skotte et al. 2013), in which we determined the best K by performing 10 replicate runs at Ks 1-7, then using CLUMPAK (Kopelman et al. 2015) to identify the best value of K. To perform both PCANGSD and NGSADMIX analyses with only *M. glaucescens* and *M. guttatus*, we generated a genotype likelihood file to include sites which contained data for 31 (out of a total of 34) individuals, a minimum mapping quality of 30, and a minimum base quality of 20, which resulted in a final dataset including 13,827,721 sites. In both cases, K=3 was optimal, so we restricted subsequent analyses to the results from PCANGSD.

#### Divergence, differentiation, and diversity

We used *Pixy* to calculate divergence, differentiation, and diversity (D_XY_, F_ST_, and π, respectively; Korunes & Samuk 2021) on 4-fold degenerate synonymous sites for all 11 taxa included herein (Northern and Southern *M. guttatus* clades were treated as distinct taxa for these analyses).For this analysis, the VCF described above was curated to include only 4-fold degenerate synonymous sites (from Brandvain et al. 2014, Coughlan et al. 2020), for the 14 main scaffolds of the *M. guttatus* reference genome (corresponding to the 14 chromosomes in this group). We used the default filtering expressions in *Pixy* (DP>=10, RGQ>=20 for invariant sites; DP>=10, GQ>=20, RGQ>=20 for variant sites), and calculated D_XY_, F_ST_, and π in 500KB windows before averaging all windows for genome-wide values (Table S5).

#### Evidence for introgression

We used Dsuite to assess Patterson’s D (i.e. ABBA-BABA statistics; Malinsky et al. 2021) for all possible trios of all individual genomes of Northern *M. guttatus* (see Results for distinction of Northern *M. guttatus* versus Southern *M. guttatus*) and *M. glaucescens*, given the ML phylogeny produced using IQtree. Our aims in this analysis were first, to assess evidence for introgression between Northern *M. guttatus* and *M. glaucescens*. Second, if significant introgression was inferred, we sought to compare the magnitude of introgression between sympatric accessions of *M. glaucescens* and *M. guttatus*, vs. allopatric accessions. Greater introgression between sympatric samples would be indicative of ongoing gene flow, as opposed to historic gene flow or more complex patterns of species bifurcation. To assess introgression we used a standard Jack-knifing procedure on all Patterson’s D statistics, with Bonferroni correction for multiple comparisons to determine if any pairs of *M. guttatus* and *M. glaucescens* genomes exhibited significant introgression. To assess whether introgression was stronger between sympatric accessions, we constructed a linear mixed model with *f*_G_ (the proportion of the genome that has experienced introgression; Malinksy *et al*. 2021) as the response variable, distribution type (whether *M. guttatus* sample was sympatric or allopatric) as fixed effect, and the individual identities of P1 in each trio as random effects. We assessed significance of the fixed effect (i.e. distribution type) with an ANOVA using a Wald’s χ^2^.

## RESULTS

### Species distribution modeling

The predicted distribution of *M. glaucescens* was centered on an area spanning the boundary of Butte and Tehama counties, whereas *M. guttatus* distribution was predicted to occur more broadly across the study area (Fig. 1A). The first principal component of the 22 modelled environmental variables explained over half of the variation observed and was positively associated with elevation and bioclimatic variables indicative of precipitation during dry and warm seasons, and negatively associated with bioclimatic variables explaining variation in temperature. PC 2 explained nearly 20% of the observed variation and was positively associated with temperature range variables and negatively associated with several precipitation variables (Fig. 1C). *Mimulus guttatus* and *M. glaucescens* differed in their mean position along PC1 (*t* = −5.93, *df* = 246.5, *P* < 0.001) for predicted occurrence points, but not PC2 (*t* = −0.407, *df* = 279.8, *P* = 0.68).

### Reproductive isolation

With the exception of ecogeographic isolation, the barriers to reproduction measured provided modest or no contribution to reproductive isolation between species (Fig. 2, Supplementary Methods and Results). For both species, the confidence intervals of reproductive isolation overlapped zero for a majority of potential barriers (Table S3), and two barriers for each species were significantly negative, indicating that hybrid matings outperformed conspecific ones for these barriers. Total isolation was estimated to be incomplete for both species, but higher for *M. glaucescens* (Fig. 2, *M. glaucescens* total RI 95% CI = 0.350-0.913; *M. guttatus* = 0.024-0.755), suggesting asymmetry in the strength of isolation. Other than ecogeographic isolation, flowering phenology (both pre- and post-zygotic) was the only barrier that was significantly positive for both species (Fig. 2). Between-population estimates of reproductive isolation within each species were also weak or overlapping zero, and broadly similar to between-species estimates (Table S3).

**Fig. 2:**
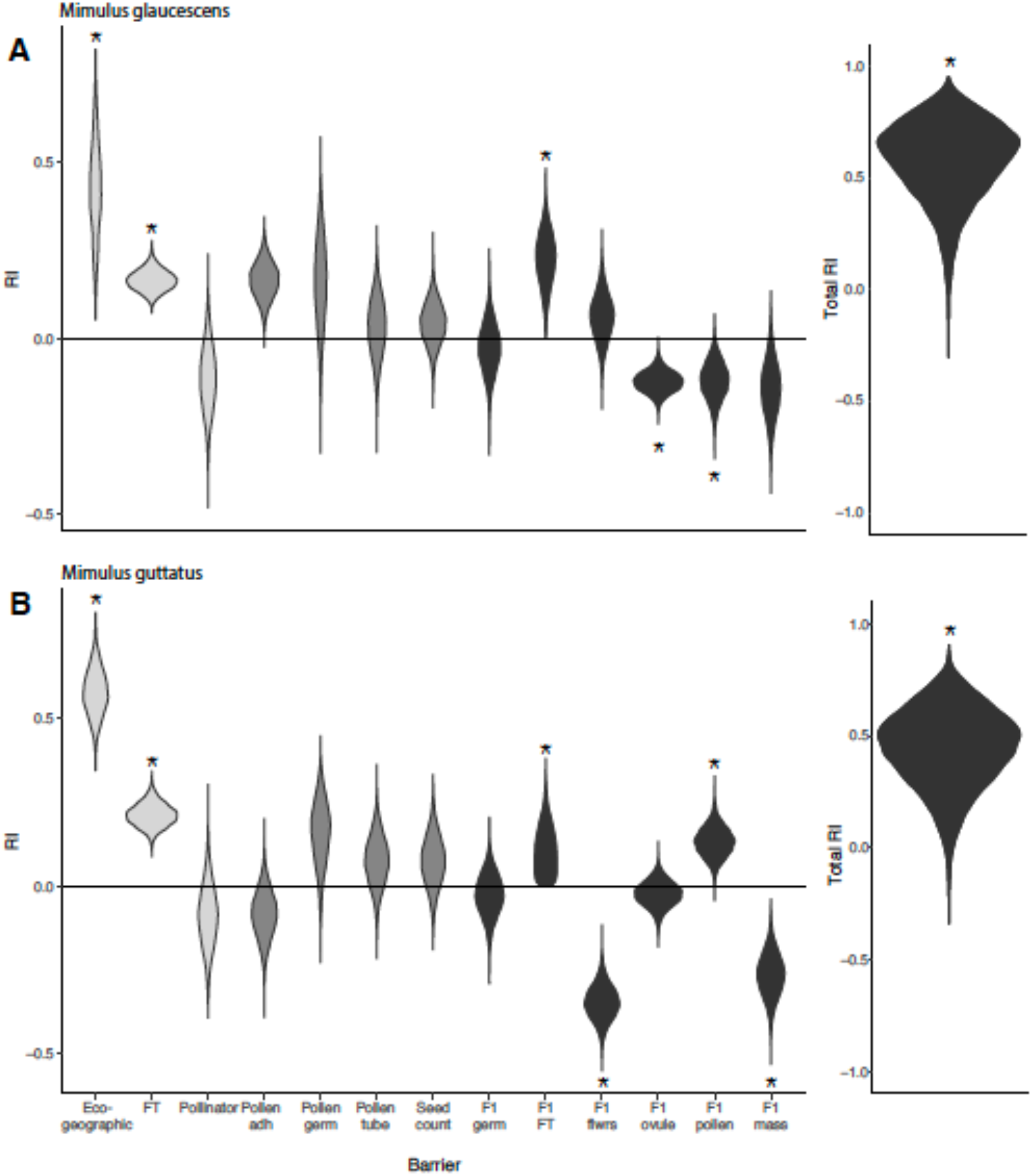
Reproductive isolation for *M. glaucescens* (A) and *M. guttatus* (B). Distributions based on 10,000 bootstrapped values (ecogeographic is based on 1000). Shadings represent types of barriers pale gray = premating barriers, dark gray = postmating, prezygotic barriers, black = postzygotic barriers. Asterisks indicate estimates with 95% CI that do not overlap 0 (Table S5).

### Genomic analyses

The complex phylogenetic history within *Simiolus* section of *Mimulus* reported elsewhere (Swegart et al. 2008; Brandvain et al. 2014; Twyford & Friedman 2015; Coughlan 2020; Coughlan 2021) was re-confirmed (Fig. 3A). The widespread taxon *M. guttatus* included deep genetic structure distinguishing a Northern and Southern clade, as has been reported elsewhere (Brandvain et al. 2014; Twyford & Friedman 2015), whereas most other named taxa included in the phylogeny represented monophyletic clades sister to one or both of the two *M. guttatus* clades. This was also the case for *M. glaucescens*, which emerged as monophyletic and sister to the Northern *M. guttatus* clade (Fig. 3A), although that monophyly was not reciprocal for Northern *M. guttatus* (similar patterns appear elsewhere within the phylogeny: Fig. 3A). Moreover, *M. glaucescens* appears to be one of the most recently diverged taxa, with both a shorter branch length and relatively lower F_ST_ and D_XY_ compared to other species pairs in the complex (Fig. 3C).

**Figure 3.**
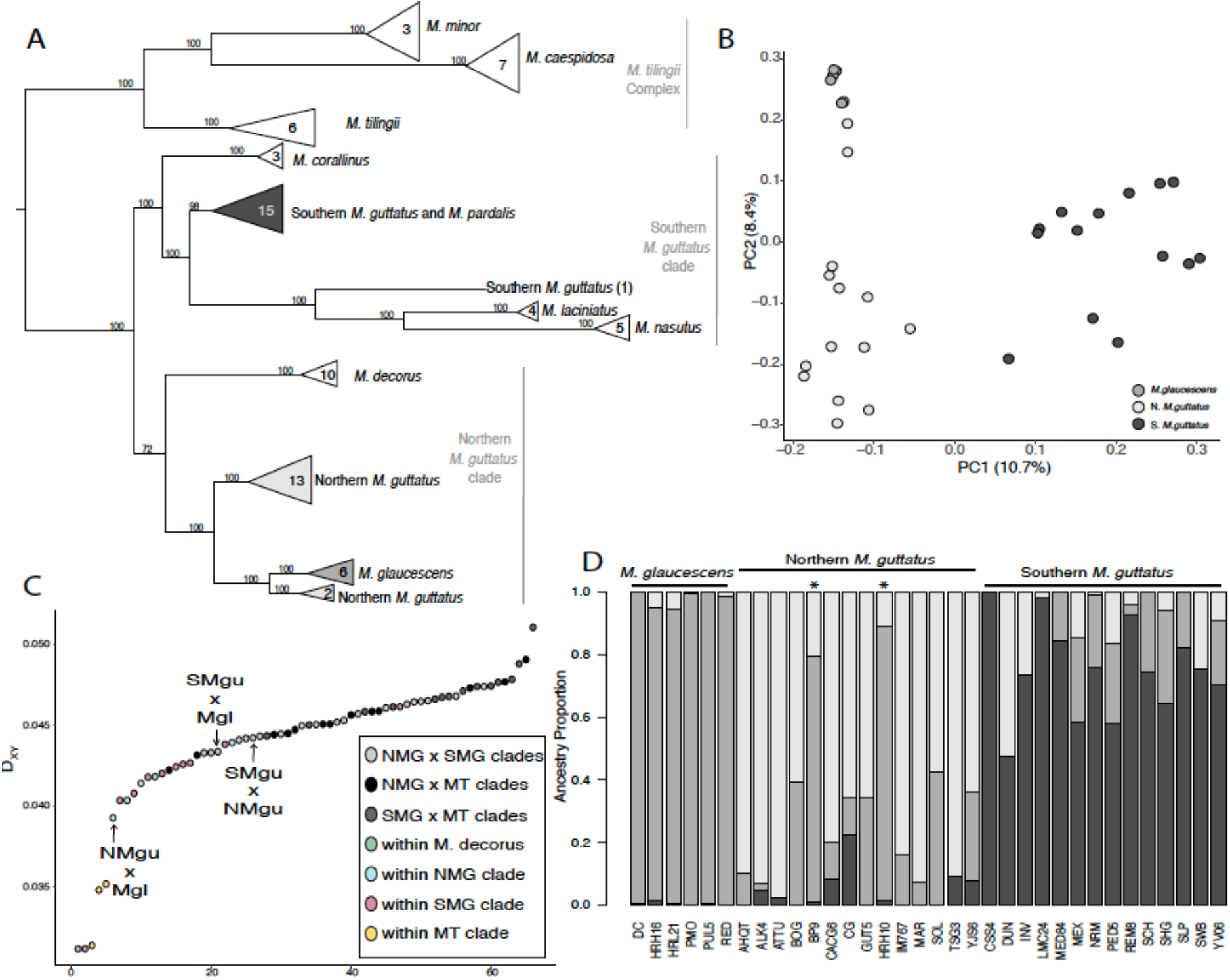
(A) Maximum-likelihood phylogeny of sampled *Mimulus* taxa based on 75 whole-genome sequences, with *M. dentilobus* (not shown) as an outgroup. Number of accessions is shown at tips next to each taxon name. (B) First two principal components of genomic variation in *Mimulus guttatus* (Southern and Northern clades represented in black and pale gray, respectively) and *M. glaucescens* (dark gray). (C) Divergence (D_XY_) in genomic variation in pairs of accessions of *Mimulus*, arranged in increasing order of D_XY_. Mean D_XY_ between selected taxa indicated by arrows (see also Table S4). NMG = Northern *M. guttatus*; SMG = Southern *M. guttatus*; MT = *M. tillingii*; Mgl = *M. glaucescens*. (D) Proportion of genome associated with ancestry of *M. glaucescens* (dark gray), Northern *M. guttatus* (light gray), and Southern *M. guttatus* (black) taxa for accessions sampled in study (see also Table S3), based on a K-means clustering analysis (see text). Asterisks indicate *M. guttatus* samples collected from within an area of sympatry with *M. glaucescens*.

The first principal component of the genome-wide data within *M. glaucescens* and *M. guttatus* mirrored the phylogenetic structure for Northern and Southern *M. guttatus* clades, in that the accessions for the two clades clustered on different ends of that axis (Fig. 3B). The second principal component, however, revealed distinct ancestry between *M. glaucescens* from *M. guttatus*, with two Northern *M. guttatus* accessions clustering with all *M. glaucescens* genomes. These two accessions had a majority (78% and 87%) of genomic ancestry characteristic of *M. glaucescens* (Fig. 3D), and were from two populations within the Butte Creek canyon area of local sympatry (Table S1). Although the ancestry percentages in these two accessions were high and were responsible for breaking reciprocal monophyly between Northern *M. glaucescens* and *M. guttatus* (Fig. 3A), *M. glaucescens*-like ancestry was relatively common across *M. guttatus* accessions (mean *M. glaucescens*-like ancestry was 24% and 10% for the Northern and Southern clade of *M. guttatus*, respectively; Fig. 3D). The reverse, however, was not observed; *M. guttatus*-like ancestry was far lower within *M. glaucescens* accessions (chiefly within sympatric accessions), consistent with the asymmetry in reproductive isolation observed (Fig. 2).

Several lines of evidence suggested recent or ongoing introgression between *M. glaucescens* and *M. guttatus. Mimulus glaucescens* and *M. guttatus* shared unusually low differentiation and divergence compared to other pairs of taxa sampled (Fig. 3C, Table S5; D_XY_ = 0.039 vs 0.043 for Northern vs. Southern *M. guttatus* clades, respectively). This result, however, varied with distribution; differentiation and divergence were higher between allopatric accessions of Northern *M. guttatus* and *M. glaucescens* vs. sympatric accessions (allopatric: D_XY_ = 0.0399, F_ST_ = 0.1090; sympatric: D_XY_ = 0.0351, F_ST_ = 0.0037), suggesting a history of introgression in sympatry. Similarly, Patterson’s D revealed significantly higher introgression between sympatric accessions than allopatric accessions of *M. glaucescens* and Northern *M. guttatus* (Fig. 4). Overall, introgression appeared to be common between both clades of *M. guttatus* and *M. glaucescens*, particularly for Northern *M. guttatus* in close proximity to *M. glaucescens*.

**Fig. 4.**
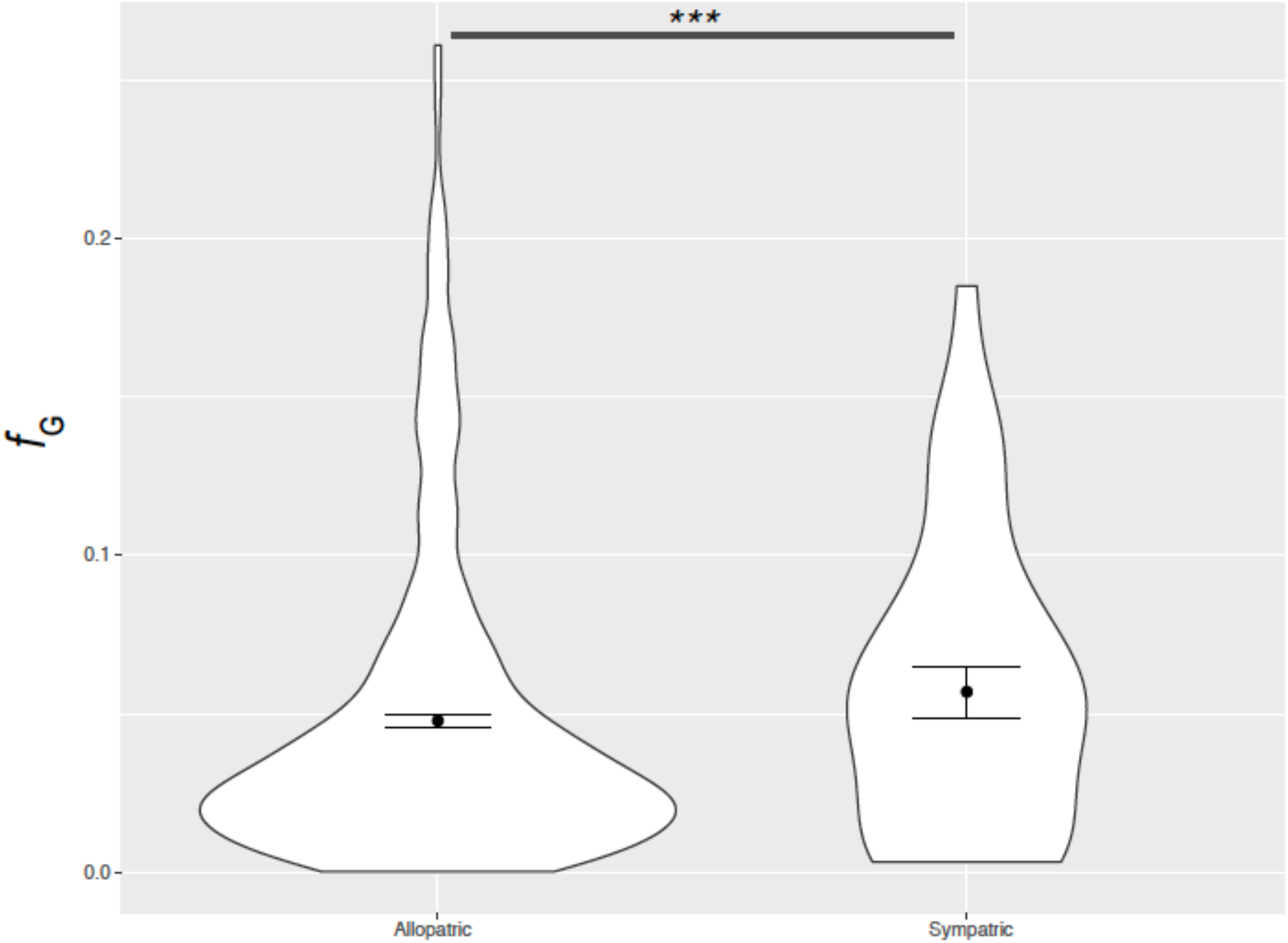
Violin plots of jack-knifed estimates of fraction of admixture (*f*_G_) in *M. glaucescens* and Northern *M. guttatus* accessions with allopatric or sympatric distributions. Estimates of *f*_G_ were significantly higher from sympatric accessions (*χ*^*2*^=91.2, *df*=1, *p*<0.0001).

## DISCUSSION

Speciation presupposes robust barriers to gene flow (Mayr 1942; Coyne and Orr 2004), thus well-defined species with less than robust isolation provoke consternation. The taxa considered herein, *Mimulus guttatus* and *M. glaucescens*, have distinctive vegetative morphology (Fig. 1B) that has led generations of taxonomists (Greene 1885; Grant 1924; Hickman 1993; Baldwin et al. 2012; Barker et al. 2012; Nesom 2012) to place them into separate species, despite nomenclatural instability in close relatives (e.g., Barker et al. 2012; Nesom 2012), and even within *M. guttatus* (Grant 1924; Nesom 2012). The morphological traits distinguishing *M. glaucescens* from *M. guttatus* are not reported as intermediate forms in the field (C.T. Ivey, personal observation) or herbaria (Nesom 2012, but see Vickery 1964). Despite divergent phenotypes, our estimates of reproductive isolation between these taxa were low (Fig. 2), with only a couple of barriers estimated to be stronger between species than between populations within a species (Table S3). Genomic evidence underscored the idea that species boundaries were porous. Allopatric accessions had greater genetic divergence, but regardless of distribution, weak genetic differentiation and significant introgression was observed between *M. glaucescens* and *M. guttatus* (Fig. 4). Overall, *M. glaucescens* should perhaps be regarded as an incipient species or an ecotype of *M. guttatus*, despite over a century of stable taxonomic status as a distinct species.

The notion that reproductively isolated or otherwise genetically distinct taxa could experience routine gene flow was historically controversial but has been widely reported more recently (Mebert 2008; Fitzpatrick et al. 2008; Martin et al. 2013; Harrison and Larson 2014; Gao et al. 2020; Jiao and Yang 2020). Persistent gene flow despite significant reproductive barriers has been observed in diverse taxa (Fitzpatrick et al. 2008; Larson et al. 2014; Richards et al. 2018; Schield et al. 2019), including other monkeyflowers (Brandvain et al. 2014; Kenny and Sweigart 2016; Nelson et al. 2021; Kiang and Hamrick 1978; Martin and Willis 2007; Case and Willis 2008; Fishman et al. 2014; Kenny and Sweigart 2016; Schemske and Bradshaw 1999; Ramsey et al. 2003). Speciation has been shown to proceed in the face of gene flow in some cases (Martin et al. 2013; Harrison and Larson 2014). Our results are thus consistent with an emerging understanding of speciation that admits that gene flow can be a routine occurrence during speciation, particularly during its early stages.

Of course, reproductive barriers limit introgression for many taxa. Prezygotic barriers are often considered to evolve more rapidly (Widmer et al. 2009; Baack et al. 2015; Christie et al. 2022), but postzygotic reproductive isolation can also be important (DiVittorio et al. 2020; Coughlan and Matutue 2020; Sandstedt et al. 2021). Regardless, the environment often mediates isolation, as with immigrant inviability (Nosil et al. 2005; Richards and Ortiz-Barrientos 2016) or selection against ecologically-unfit hybrids (Jacquemyn et al. 2018; Ferris and Willis 2018). For *M. glaucescens* and *M. guttatus*, the strongest barrier was ecogeographic, with climate and elevation as potential agents of ecological differentiation (Fig. 1). Ecogeographic isolation is often strong in plants (Lowry et al. 2008a, 2008b; Sobel 2014; Li et al. 2018; Christie & Strauss 2018; Vargas et al. 2020; Sianta and Kay 2021), although divergence in habitat is often associated with genetically based differences in reproductive biology, such as flowering time (Silvertown et al. 2005; Antonovics 2006; Bomblies 2010; Sianta and Kay 2021) or pollination (Grossenbacher and Whittall 2011; Ostevik et al. 2012). Otherwise, assortative mating within taxa may be limited, with free genomic exchange, except for loci explicitly involved in local adaptation (e.g., Hendrick et al. 2016). In addition to ecogeographic isolation, we found significantly positive, though weak, isolation based on divergence in flowering time (Fig. 2), which is consistent with the idea that habitat and reproductive divergence may co-occur during speciation. Despite weak isolation, the distinctive genetic coherence of *M. glaucescens* was supported based on the phylogenetic analysis (Fig. 3) as well as estimates of differentiation; for example, D_XY_ between Northern *M. guttatus* and *M. glauscens* was higher than π (nucleotide diversity) within Northern *M. guttatus* (Fig. 3; Table S4).

The factors maintaining vegetative differences between *M. glaucescens* and *M. guttatus* (Fig. 1) remain unclear. To our knowledge the genetic basis for expression of these traits (peduncular bract fusion and glaucousness, trichomes) in *M. glaucescens* has not been characterized, but large-effect QTL for glaucousness and trichome density reported from other studies (Bennet et al. 2012; Holeski et al. 2010; Hendrick et al. 2016) suggest that segregating phenotypes might not be intermediate. If so, hybrid or introgressed individuals may be difficult to discern in the field based on vegetative phenotype alone. Glaucousness has been found to facilitate drought tolerance (Xue et al. 2017) and defense against insect herbivores (Eigenbrode and Espelie 1995). Similarly, transitions in trichome density have been shown to be influenced by natural selection within other monkeyflower taxa (Hendrick et al. 2016). The extent to which these traits contribute to the divergence observed in habitat distribution between *M. glaucescens* and *M. guttatus* remains to be tested.

The pattern of distribution observed – asymmetric range sizes, broadly sympatric, but with environmentally based geographic divergence – was similar to that reported for other pairs of *Mimulus* sister taxa (Grossenbacher et al. 2014; Sobel 2014) and indeed, pairs of sister taxa more generally within the California Floristic Province (Anacker and Strauss 2014). This pattern is not inconsistent with models of budding (or peripatric) speciation (Grossenbacher et al. 2014). The genomic analyses are also consistent with the idea that budding speciation is common throughout the species complex; many of the taxa included in the phylogeny, including *M. glaucescens*, were sister to one of the main *M. guttatus* clades (Fig. 3A). Similarly, *M. guttatus* ancestral variation was substantially represented within each of the putatively derived taxa (Fig. 3A; Fig. S1; Table S5).

In conclusion, we show that the recently diverged, but well differentiated taxa, *M. glaucescens* and *M. guttatus*, have surprisingly low reproductive isolation and abundant gene flow, in contrast to what is suggested by their taxonomy and morphology. This study contributes to a growing understanding of the complex evolutionary patterns involved in the formation of species boundaries during speciation (Martin et al. 2013; Feder et al. 2014; Harrison and Larson 2014; Jiao and Yang 2020; Stankowski and Ravinet 2021; Coughlan et al. 2021) and the contribution of reproductive barriers toward delineation of lineages (Baack et al. 2015; Ostevik et al. 2016; Christie and Strauss 2019; Runemark et al. 2019). More fundamentally, these results underscore that integration of classical ecological studies of reproductive isolation with genomic approaches will be instrumental for continued progress in disentangling this fundamental and intriguing evolutionary process.

## Supporting information

Supplemental methods & results

## ACKNOWLEDGEMENTS

We acknowledge the indigenous native peoples of California as the traditional stewards of the ecosystems studied for this project. K. M. Kay, J. M. Sobel, and the A. L. Sweigart lab commented on the manuscript; L. S. Kimsey of the University of California, Davis Bohart Museum identified floral visitors; L. Bergmann, C. Bruns, T. Devine, J. Habecker, M. Patterson, and A. Pokrzywinski helped in the field and greenhouse; K. A. Blee, K. F. Gorman, C. Hatfield, J. M. Sobel, D. G. Miller, and K. A. Schierenbeck provided advice and suggestions; and the Big Chico Creek Ecological Reserve, California Botanical Society, California Native Plant Society, Chico State Center for Water and the Environment, Friends of the Chico State Herbarium, Northern California Botanists, and the Vesta Holt Field Research Award provided financial support.

