## Supplemental methods & results for "When good species have porous boundaries: weak reproductive isolation and extensive gene flow between *Mimulus glaucescens* and *M. guttatus* in northern California"

Supplementary Methods and Results

### METHODS

#### Species distribution modelling

To estimate the potential for broad-scale divergence in habitat to limit interspecific reproduction, species distribution models were constructed based on climate, soils, and elevation. Primary and secondary geological substrates (Goff et al. 2021), elevation (National Elevation Datatset: http://ned.usgs.gov/), and 19 Bioclimatic variables (https://www.worldclim.org/) from within the study area were all rasterized to a standard resolution of 2.5 min. In addition, georeferenced collection records from the Consortium of California Herbaria (https://ucjeps.berkeley.edu/consortium/), the Global Biodiversity Information Facility (http://www.gbif.org/, excluding iNaturalist observations), and Mimubase (http://mimubase.org/accessions) were compiled. Pseudo-absence data were generated by randomly sampling 1000 background sites, bounded by -121 and -123 longitude and 39 and 40.5 latitude (approximately the extent of Butte and Tehama counties, where *M. glaucescens* occurs).

The environmental and occurrence data were combined using the *raster* (Hijmans *et al.* 2012; Hijmans 2020) and *dismo* packages in R (Hijmans *et al.* 2017). We then constructed maximum entropy species distribution models using the *maxnet* function in the *dismo* package in R (Hijmans *et al.* 2017) for each species separately, using 75% of the data randomly selected to train the model and the remaining 25% to test its performance, iterated for 1000 replicates. The threshold for determining occurrence of a species within a location was determined using the area under the receiving operator characteristic curves (Phillips et al. 2006), while balancing sensitivity with specificity by using the "spec_sens" statistic under the *threshold* function in *dismo*.

To assess differences in species distribution, we first used a principal components analysis from the predicted occurrence locales with all environmental variables included. PCs 1-3 explained significantly more variation than expected under a broken stick model. As PCs 1 and 2 explained the vast majority of variance (54.4% and 19.6%, respectively), We performed a Welch's *t-test* separately for PC1 and PC2 to determine if *M. glaucescens* and *M. guttatus* differed significantly in habitat.

To characterize isolation based on habitat, the number of pixels on the final distribution map that were predicted to be occupied by each species separately as well as both species together was calculated. Areas of predicted shared habitat were considered to have a higher chance of interspecific gene flow than areas of predicted unshared habitat. The number of shared and unshared pixels was used to estimate $RI_{ecogeographic}$ using equation $RI_{4C}$ from Sobel and Chen (2014). Note that *M. guttatus*, because of its broad western North American distribution, is geographically isolated from *M. glaucescens* throughout most of its range. Thus, estimates of $RI_{ecogeographic}$ for *M. guttatus* based on these models would underestimate species-level isolation and would infer isolation only for populations within the region modelled.

#### Habitat in sympatry

In addition to estimating broader geographic patterns of habitat differentiation above, we collected local-scale environmental data in a 6 $km^2$ area of sympatry (Butte Creek canyon, Butte County) to characterize habitat differentiation at a finer scale, which also may contribute to reproductive isolation (*sensu* Li et al. 2018). At each of nine populations of *M. glaucescens* and six populations of *M. guttatus*, 8-10 plants were randomly chosen. For each plant, slope was estimated by placing a clinometer (Suunto Tandem, Vantaa, Finland) at its base, parallel to the main slope axis, and recording angle in degrees. Percent shade cover was estimated for each plant as the average of four readings, in each of the cardinal directions, of a spherical densiometer (Forest Densiometers, Bartlesville, Oklahoma) held level at the apex of each plant. In addition, a 0.25 $m^2$ quadrat was centred on each plant and used to estimate percent vegetative cover, percent bare ground, percent debris, and percent standing water. All measurements ($n$ = 144 plants) were standardized to mean = 0 and SD = 1 before conducting a principal components analysis using the six habitat variables. Scores of the first two principal components were compared using a mixed-model ANOVA with species as the fixed effect and population as the random effect. Significant differences between species were interpreted as evidence for habitat segregation. All statistical analyses were conducted using SAS 9.3 (SAS Institute, Cary, NC), unless otherwise indicated.

Flowering phenology

To estimate the contribution of flowering phenology to temporal reproductive isolation, eight populations (four of each species) co-occurring within a 2 $km^2$ area of Butte Creek canyon were monitored for one flowering season (Table S1). Populations were placed into pairs within a species as well as between species based on proximity, elevation, and size. Pairs were separated by a mean (SD) of 1.6 (1.2) km, which is within the flight range of the predominant floral visitors (Beekman and Ratnieks 2000, Gathmann and Tscharntke 2002, Wolf and Moritz 2008; Guédot et al. 2009; Table S2). All populations occurred in moist areas bordering a roadway. At each site the number of flowering plants was recorded once each week from mid-March (when no plants were flowering) until early August (after all flowering had stopped). Reproductive isolation due to phenology ($RI_{phenology}$) was calculated using equation $RI_{4S2}$ from Sobel and Chen (2014), which incorporates information about relative abundance as well as differences in flowering time. Estimates of isolation were calculated for pairs of populations within species as well as between species. $RI_{phenology}$ between populations within species was calculated for each within-species population pair (in both directions), and averaged within species to estimate between-population isolation for each species. To estimate isolation between species, $RI_{phenology}$ was calculated for each species (i.e., in both directions) within each between-species pair of populations, and then averaged across all pairs within each species for an overall estimate. In addition, we compared flowering plant numbers between species using a repeated measures model in which plant species, day of observation, and their interaction were fixed effects, and the between-species pair identity was a random effect. The day of observation was the repeated effect with the population as the subject. Flowering plant number was natural log-transformed prior to analysis to correct heteroskedasticity, but model results are presented on the original scale.

Pollinator visitation

To estimate reproductive isolation due to pollinator behavior, floral visitors were observed in experimental mixed-species arrays near natural populations of each species. Arrays were placed near existing populations in flower to ensure the presence of naturally occurring pollinators. One set of arrays was within the Big Chico Creek Ecological Reserve (BCCER) near a population of *M. glaucescens*, and a second in Butte Creek canyon near a population of *M.*

*guttatus* (Table S1). Each array consisted of 18 flowering plants of each species (36 total) collected from nearby natural populations and transplanted into 9 $cm^2$ pots. Pots were arranged into a 6 x 6 checkerboard grid, at 0.5 m spacing, which was comparable to the density of nearby populations. Floral visitors would thus have an equal chance of encountering either species when moving to a neighboring plant. To maintain equivalence in floral display size, corollas were removed from older open flowers if necessary such that each plant displayed one open flower, each of which had opened that day. Arrays were observed in 1-hr observation periods from 0900 to 1500 during the two weeks of peak flowering (total 67 hr). New arrays were assembled for each day of observation; during each day, multiple observations were collected from an array. For each floral visitor, the taxon and the sequence of plants visited ("observation") was recorded. For observations that involved visits to > 1 plant, the likelihood of conspecific (assigned a value of 0) vs. heterospecific (1) movements was compared using a generalized linear mixed model to which a binomial error distribution was assigned. Fixed effects included the plant species from which the visitor moved, site location, and their interaction. Random effects included visitor taxon (Table S2) and the observation. A probability of heterospecific movement significantly < 0.5 was considered to be evidence of pollinator preference, which may contribute to reproductive isolation. Reproductive isolation due to pollinator preference ($RI_{pollinator}$) was calculated using equation $RI_{4A}$ from Sobel and Chen (2014). A significant difference between plant species in the likelihood of visitors switching species (i.e., heterospecific movements) was considered to be evidence of asymmetry in pollinator preference.

Flower morphology

To estimate indirectly the extent to which species boundaries might be maintained by differences in floral morphology (through mechanical isolation, *sensu* Gögler et al. 2015, or other mechanisms), flower morphology was compared between species. Floral traits have not been used to distinguish these taxa (e.g., Baldwin et al. 2012; Nesom 2012), but to our knowledge their flower morphology had not been compared closely before. Corolla width at the widest point, corolla tube width at distal opening, anther length, pistil length, and herkogamy were measured to the closest mm for 120 first-day flowers (*M. glaucescens* $n$ = 73, mean [SD] plants per population = 18.0 [9.1]; *M. guttatus* $n$ = 47, mean [SD] plants per population = 11.8 [10.9]) using plants reared from open-pollinated seeds collected in four populations of each species (Table S1). Plants were grown in a greenhouse in 25 $cm^2$ pots filled with Promix HP (Premier Tech Horticulture, Rivière-du-Loup, Quebec, Canada) soilless media. Pots were arranged randomly within trays, provided with water ad libitum, fertilized three weeks after sowing with 800 ml of Peter's 20-20-20 (Scotts, Marysville, Ohio, USA) at 1.5 g $L^{-1}$, and provided with 18 hr daily supplemental lighting from 400W sodium vapor lamps. Family mean values of trait measurements were standardized to mean = 0 and SD = 1 and a principal components analysis was applied to the data. Scores of the first two principal components were compared between species using a mixed-model ANOVA with species as the fixed effect and population as a random effect. A significant difference between species was considered evidence of divergent floral morphology.

Post-pollination pollen-pistil interactions

Pollen adhesion, pollen germination, and pollen tube growth rate (PTGR) were compared between conspecific and hybrid hand-pollinations using a semi *in vitro* experimental approach to estimate their potential for limiting the likelihood of gene exchange (*sensu* Kay 2006). For the adhesion and germination experiments, plants were grown as above from open-pollinated seeds collected from four populations (two of each species) within Butte Creek canyon (Table S1). Additional populations were included for the PTGR experiment (Table S1). Flowers were randomly assigned as pollen recipients and emasculated in bud one day prior to anthesis to prevent self-pollination. Pollen donors were matched randomly with recipients.

*Adhesion* - Differences between species in pollen adhesion to stigmas could contribute to reproductive isolation by affecting the success of pollination involving mixed or heterospecific deposition. Methods modified from Zinkl et al. (1999) were used to characterize pollen adhesion. Freshly dehisced anthers from a randomly chosen pollen donor were gently tapped to sprinkle pollen onto a clean microscope slide. Under a dissection microscope at 40x, pollen was then removed from the slide until approximately 100 grains remained, after which the number of pollen grains was counted. The pistil of a previously emasculated, first-day flower randomly chosen as pollen recipient (sample size *M. glaucescens* within-population = 11, between-population conspecific = 6, heterospecific = 25; *M. guttatus* = within-population = 16, between-population conspecific = 9, heterospecific = 28) was then removed from the flower and the pollen grains were wiped from the slide using the receptive stigmatic surface. Any pollen grains remaining on the slide were counted and subtracted from the original number of grains on the slide to calculate the number of pollen grains deposited onto the stigmatic surface. After 60 sec pistils were placed into a microcentrifuge tube with 200 µl of 50µM $KPO_4$ with 1% Tween-20, vortexed for 5 seconds, centrifuged for 60 sec at 7200 rpm, and then fixed in a solution of 9:1 ethanol:glacial acetic acid for 24 hr. Fixed stigmas were stained with basic fuchsin dye and viewed at 100x to record number of pollen grains adhering to stigmatic surfaces. Proportion of pollen adhering was arcsine square-root transformed prior to analysis to correct heteroskedasticity, but results are reported on the original scale.

*Germination* - To examine the potential for variation in pollen germination to contribute to reproductive barriers, the germination of pollen following hand-pollinations was compared among cross types. Clean micro-forceps were inserted roughly 1 mm into a freshly dehisced anther of a randomly chosen pollen donor. The pollen-coated forceps were then placed between lobes of randomly chosen recipient stigmas, which resulted in a mean (CV) of 312 (0.76) pollen grains transferred to stigmas (sample size *M. glaucescens* within-population = 15, between-population conspecific = 20, heterospecific = 26; *M. guttatus* = within-population = 15, between-population conspecific = 10, heterospecific = 31). After sufficient time for germination to commence (6-12 hours), pistils were harvested and fixed in a 3:1 solution of 95% ethanol and glacial acetic acid for 24 hours. Pistils were rinsed with deionized water and stored in 70% ethanol until observation. After clearing in 70°- 80°C 1 M NaOH for 2 minutes, pistils were rinsed in deionized water and stained in decolorized 0.1% aniline blue in 0.033 M $K_3PO_4$ for 24 hours. Pistils were then mounted on slides with a drop of aniline blue, gently squashed with a coverslip, and viewed at 100x magnification under epifluorescence (Leitz Ortholux II, Wetzlar, Germany). The number of pollen tubes within styles and the number of pollen grains on stigmas was recorded. Proportion of pollen grains germinated was analyzed on the original scale.

*Pollen tube growth rate* - Pollen tube growth rate was evaluated for its potential to contribute to reproductive isolation. To perform pollinations, a single freshly dehisced anther from a randomly chosen pollen donor was gently pressed into the stigma of a randomly chosen recipient flower (sample size *M. glaucescens* within-population = 7, between-population conspecific = 11, heterospecific = 17; *M. guttatus* = within-population = 9, between-population conspecific = 12, heterospecific = 21). After five minutes, styles were cut cleanly from pistils with stigmas attached, and placed on 1% agar enriched with 0.3 M sucrose, 1.6 mM $H_3BO_3$, 2 mM $CaCl_2$, and 1 mM $MgSO_4 \cdot 7H_2O$. Style length was measured to the nearest 0.1 mm using a dissection microscope fitted with an ocular micrometer. The time in min until pollen tubes were observed emerging from the cut end of the style was recorded. Rate of pollen tube growth was then calculated as mm/min, and it was analyzed on the original scale.

*Analysis of pollen-pistil interactions* – Components of pollen-pistil interaction performance were evaluated using two-way mixed-model ANOVA. Model fixed effects included the category of hand-pollination ("between-population conspecific," "within-population," or "heterospecific"), the species of pollen recipient, and their interaction. Random effects included the population identities of pollen donor and pollen recipient. The magnitude of reproductive isolation due to each pollen-pistil interaction component ($RI_{adhesion}$, $RI_{germination}$, $RI_{PTGR}$) was calculated between populations within each species as well as between species using equation $RI_{4A}$ from Sobel and Chen (2014).

<u>Additional intrinsic post-pollination barriers</u>

*Experimental design* – To characterize other components of intrinsic post-pollination performance, one randomly chosen seedling from each of 100 open-pollinated families from each of two populations each of *M. glaucescens* and *M. guttatus* (Table S1) was transplanted into 25 $cm^2$ pots with equal parts peat, sand, pumice, and redwood compost, and provided with 12-hr supplemental fluorescent light (six-tube fixtures at 40 cm with 6500 K, 2850 lumen tubes) at room temperature. Plants were watered as needed and fertilized every 14 d with 16-16-16 NPK fertilizer (MaxSea, Garberville, CA). After 40 days, temperature was lowered to 4°C for 32 days for vernalization, after which it was returned to 22°C, and photoperiod increased by 30 minutes every 2-3 days until reaching 16 hr. Flowering plants were assigned at random to serve as pollen donor or recipient. Flowers assigned as pollen recipients were emasculated in bud a day before anthesis. A single freshly dehisced anther from a plant assigned as pollen donor was gently pressed into the stigma lobes of its assigned recipient to pollinate the flower.

*Seed production* – Fruits (sample size *M. glaucescens* within-population = 41, between-population conspecific = 20, heterospecific = 58; *M. guttatus* = within-population = 63, between-population conspecific = 32, heterospecific = 90) were collected as they matured and seeds from each photographed digitally on white paper. Images were imported to ImageJ (Schneider et al. 2012) to count seeds following Ivey and Carr (2012). Seed number per fruit was compared among categories of crosses.

*Seed germination* – Twenty mature fruits of each category of cross (within-population, between-population conspecific, and heterospecific) from each species were randomly chosen, and 20 seeds from each were sown into 100 $cm^2$ pots under cultivation conditions described above. After two weeks, the number of seedlings was counted in each pot. Proportion of seeds germinated was compared among categories of crosses.

*Post-germination performance* – One randomly chosen seedling from each of the subset of fruits chosen for the germination experiment (2 species x 20 fruits x 3 cross categories = 120 total plants) was transplanted into a 100 $cm^2$ pot and grown to maturity as described above. For each plant, the number of days from sowing until first flower opened ("DFF") was scored as a measure of developmental rate. In addition, ovule number and pollen viability were scored for the first flower to open as estimates of female and male fertility, respectively (Carr and Dudash 1997). Harvested pistils were placed in 3:1 ethanol: glacial acetic acid until assays were performed. Ovules from a single carpel per flower were counted under a light microscope after staining with lactophenol aniline blue. All anthers from each harvested flower were placed in lactophenol in microcentrifuge tubes until assays were performed. The proportion of stained pollen grains in three 25 µl aliquots was averaged for each sample. After 59 days of growth, total number of flowers was recorded for each plant. On day 60, all above-ground biomass was harvested, dried to constant mass at 55º C, and weighed to the nearest mg.

*Analysis of post-pollination performance* – Means of post-pollination performance traits were compared using generalized linear mixed-models. Fixed effects for all models included the category of hand-pollination ("between-population conspecific," "within-population," or "heterospecific"), the species of female parent, and their interaction. Random effects included the population identities of paternal parent and female parent. A beta distribution with a logit link function was applied to seed germination and pollen viability data; a Poisson distribution with a log link function was applied to seed number, DFF, flower number, and biomass; and a Gaussian distribution with an identity link function was applied to ovule number. Reproductive isolation for each component was calculated between populations within each species as well as between species using equation $RI_{4A}$ of Sobel and Chen (2014), with the exception of DFF, for which equation $RI_{4C}$ was used to evaluate overlap in flowering.

Genomic analyses:

We added to previously available sequence data by sequencing an additional 11 genomes. First, genomic DNA was extracted from fresh or silica-gel dried leaf and/or bud tissue using a modified CTAB protocol (Kelly & Willis 1998; dx.doi.org/10.17504/protocols.io.bgv6jw9e) or using a Qiagen DNeasy Plant Pro kit, after which libraries were prepared by the Duke Sequencing facility, the University of California Davis Genome Center, or by using Illumina Nextera DNA (as in Puzey et al. 2018; Coughlan et al. 2020; Coughlan et al. 2021). Individually barcoded libraries were then pooled in equal molar amounts, and were sequenced using either a HiSeq 2500 Rapid-Run or NovaSeq S4 platform, to produce 150bp, paired end reads with 5.8x average read depth (see Table S4 for details).

To process the resultant reads, we trimmed adapters and low-quality sequences using TrimGalore! (Krueger 2014), aligned reads to the *M. guttatus* V2.0 hardmasked reference genome using the BWA *mem* function (Li et al. 2009), and cleaned, sorted, and marked duplicates in the resulting bam files using Picard tools (http://broadinstitute.github.io/picard/). Last, we used the *HaplotypeCaller* function in GATK to identify SNPs and call genotypes for each accession individually, then the *GenotypeGVCFs* function in GATK to jointly call genotypes across all samples for both variant and invariant sites (McKenna et al. 2004). We filtered the resultant VCF file to remove INDELs, keep sites with a minimum quality score (minQ) of 30 and a minimum coverage of 5x per sample using VCFtools (Danecek et al. 2011). Subsequent filtering of the VCF file was performed to best suit each analysis, described below.

### RESULTS

Pre-pollination barriers

*Local-scale habitat* - The first principal component explained 32% of the variance in the environmental data collected, and it was most strongly correlated with slope and canopy (both positively) and vegetative cover (negatively). The second principal component explained an additional 27% of the variance, and it was most strongly correlated with debris cover, vegetative cover, and canopy cover (all positively) as well as percent bare ground (negatively). Neither the first ($F_{1,\,129}$ = 0.64, $P$ = 0.4) nor the second ($F_{1,\,129}$ = 0.52, $P$ = 0.5) principal component significantly explained habitat differences between species where they occurred in sympatry (Fig. S2).

*Flowering phenology* - Between populations of *M. glaucescens*, flowering onset was separated by an average of four weeks, and peak flowering by 2 weeks. Between populations of *M. guttatus*, flowering onset was separated by one week, and peak flowering was offset by an average of 3.5 weeks. Between species, *M. glaucescens* began flowering three and five weeks earlier than *M. guttatus* in two of the populations, whereas for the other two, flowering onset was nearly concurrent (separated by 1 week or less). Peak flowering occurred later for *M. guttattus* (mean [SD] = 15.8 [8.8] days later). The end of flowering was within two weeks (mid- to late-July) for all pairs. Populations of *M. glaucescens* had more flowering plants on average than *M. guttatus* (species effect; $F_{1,\,111}$ = 5.75, $P$ = 0.02; least-squares mean[SE] flowering plants *M. glaucescens* = 263[42], *M. guttatus* = 197[42]), but this did not depend on the day of observation (species x day interaction; $F_{18,\,111}$ = 0.99, $P$ = 0.5; Fig. S3).

*Pollinator visitation* - Of 533 flower visits recorded, over 40% (223) involved movements between plants, 101 (45.3%) of which were conspecific and 122 (54.7%) heterospecific. All species of floral visitors were observed visiting both plant species (i.e., no visitor taxa were unique to a species; Table S2). Visitors foraging on *M. guttatus* had a least-squares mean (SE) = 0.52 (0.11) frequency of moving heterospecifically, whereas those foraging on *M. glaucescens* had least-squares mean (SE) = 0.66 (0.14) frequency of heterospecific transitions, and these did not significantly differ between species ($F_{1,\,216}$ = 0.48, $P$ = 0.49), suggesting no asymmetry in preference between species. The 95% confidence intervals for both species overlapped 0.5 (*M. glaucescens* = 0.36-0.87; *M. guttatus* = 0.30-0.74), suggesting that visitor movement between species was random. A log-likelihood test of the random effect of visitor taxon suggested there were no significant differences among visitors in movement ($X^2$ = 2.53, $P$ = 0.06).

*Flower morphology* - The first principal component (PC) of the floral trait measurements explained over 50% of the variance in the data, and the factor loading scores were all positive, suggesting that these PC scores reflected variation in size among flowers. The second PC explained an additional 31% of the variance. Tube length, corolla width, and herkogamy were positively associated with the second PC, whereas pistil length and anther length were negatively associated. Neither the first ($F_{1,\,6.3}$ = 0.01, $P$ = 0.9) nor the second ($F_{1,\,44}$ = 2.10, $P$ = 0.2) PC scores differed significantly between species (Fig. S4), thus it appeared that opportunities for reproductive isolation based on flower morphology were negligible.

<u>Post-pollination barriers</u>

*Adhesion* – About 20% fewer pollen grains adhered to the stigmas of *M. guttatus* than to *M. glaucescens* (Fig. S5A, $F_{1,\,3.3}$ = 6.94, $P$ = 0.07), but adhesion did not vary among categories of experimental pollinations performed ($F_{2,\,88.3}$ = 0.12, $P$ = 0.9), nor did the pollen recipient species x pollination category interaction contribute toward explaining variation in adhesion ($F_{2,\,88.3}$ = 0.88, $P$ = 0.4).

*Germination* – The proportion of pollen grains that germinated on stigmas did not differ between recipient species (Fig. S5B, $F_{1,\,1.9}$ = 1.0, $P$ = 0.4) or, on average, among categories of pollination ($F_{2,\,103}$ = 0.24, $P$ = 0.8). The effect of pollination category on germination, however, was not equivalent between the recipient species (recipient species x pollination category interaction: $F_{2, 50.4} = 5.94$, $P = 0.005$). Specifically, germination in heterospecific pollinations of *M. guttatus* was nearly 10% lower than within-population pollinations (Fig. S5B, $P = 0.04$ in a Tukey's post-hoc test). Germination from between-population crosses of *M. guttatus* was about 3% higher than within-population crosses, but this difference was not significant ($P = 0.9$ in a Tukey's post-hoc test). Although pollen germination in heterospecific pollinations of *M. glaucescens* was about 6% higher than for within-population crosses, and nearly double that of between-population crosses (Fig. S5B), these differences were not significant ($P > 0.05$ in Tukey's post-hoc tests).

*Pollen-tube growth rate* – Pollen tubes grew about 3 µm/min faster in the styles of *M. glaucescens* flowers (Fig. S5C, $F_{1, 54} = 4.86$, $P = 0.03$) regardless of the category of pollination performed (category of pollination: $F_{2, 54} = 0.3$, $P = 0.7$; recipient species x pollination category interaction: $F_{2, 54} = 0.54$, $P = 0.6$).

*$F_1$ seed number* – Crosses in which *M. glaucescens* was the female parent produced about half the seeds of those involving *M. guttatus* as pollen recipient (Fig. S6A, $F_{1, 4} = 8.20$, $P = 0.05$). In addition, seed number depended on the type of cross performed, on average ($F_{2, 298} = 664.4$, $P < 0.0001$), although this was not equivalent between the two species (female parent x cross type interaction $F_{2, 4.7} = 14.6$, $P = 0.01$). Specifically, between-population crosses resulted in a 23% increase in seed set for *M. glaucescens*, but only a 9% increase for *M. guttatus* (Tukey's post-hoc tests $P < 0.007$). Heterospecific crosses, on the other hand, produced fewer seeds for both species, although not significantly so (Tukey's post-hoc tests > 0.05).

*Seed germination* – Proportion of seeds germinating did not differ between female parent species (Fig. S6B, $F_{1, 2} = 0.15$, $P = 0.7$) or among cross types ($F_{2, 116} = 0.31$, 0.7), nor was there a significant interaction between female parent species and cross type ($F_{2, 3.7} = 0.06$, $P = 0.9$).

*Biomass* – Final above-ground dry biomass did not differ between plants derived from the two female parental species (Fig. S6C; $F_{1, 138} = 1.44$, $P = 0.2$) or among types of crosses ($F_{2, 177} = 0.25$, $P = 0.8$), nor did final biomass depend on the interaction between female parent and cross type ($F_{2, 3.2} = 0.67$, $P = 0.6$).

*Days to first flower* – The reproductive developmental rate was about the same, on average, for plants derived from *M. glaucescens* female parents as those involving *M. guttatus* as female parent (Fig. S6D, $F_{1, 2.2} = 0.01$, $P = 0.9$). The development of between-population crosses was nearly the same as within-population crosses for both species (Tukey's post-hoc tests $P > 0.9$). Hybrid plants, however, took about 4.8 days (13%) longer to reach the flowering stage than non-hybrid plants, on average (cross type main effect: $F_{2, 152} = 9.84$, $P < 0.0001$; Tukey's post-hoc tests $P < 0.004$). Differences in development among cross types did not differ between the two species (cross type x female parent interaction: $F_{2, 4.6} = 0.13$, $P = 0.8$).

*Flower number* – Total number of flowers produced did not differ between species serving as female parent (Fig. S6E, $F_{1, 2.7} = 0.61$, $P = 0.5$), but there were differences among cross types ($F_{2, 150} = 21.6$, $P < 0.0001$), which depended on female parental species (cross type x female parent interaction: $F_{2, 4.6} = 43.5$, $P = 0.001$). Specifically, between-population as well as hybrid plants in which *M. glaucescens* served as female parent produced fewer flowers than within-population *M. glaucescens* crosses. In contrast, between-population and hybrid crosses from *M. guttatus* maternal parents produced more flowers than within-population *M. guttatus* crosses.

*Ovule number* – Carpels from flowers in which *M. glaucescens* served as female parent had about 20% fewer ovules than those in which *M. guttatus* was the maternal parent, on average (Fig. S6F, $F_{1, 16.6} = 12.7$, $P = 0.003$), but ovule number did not vary among cross types ($F_{2, 151} =$

0.85, *P* = 0.4), nor was there a significant cross type x female parent interaction ($F_{2, 4.2}$ = 0.75, *P* = 0.5). Flowers on between-population crosses from *M. glaucescens* had nearly the same number of ovules per carpel as within-population crosses, and hybrid plants produced from *M. glaucescens* maternal parents had about 7% more ovules than those from within-population crosses. Between-population crosses involving *M. guttatus*, on the other hand, had fewer ovules per carpel, as did flowers on hybrids with *M. guttatus* maternal parents.

*Pollen viability* – The proportion of viable pollen did not significantly differ on flowers derived from crosses in which *M. glaucescens* was the maternal parent vs. those in which *M. guttatus* served as female parent (Fig. S6G, $F_{1, 2.1}$ = 0.01, *P* = 0.09), nor did this vary among cross types ($F_{2, 148.6}$ = 0.26, *P* = 0.8). In addition, there was no significant cross type x female parent interaction ($F_{2, 4.5}$ = 1.66, *P* = 0.3).
