## supplemental tables and figures for "When good species have porous boundaries: weak reproductive isolation and extensive gene flow between *Mimulus glaucescens* and *M. guttatus* in northern California"

Supplementary Tables and Figures

Table S1. Locations of populations of *Mimulus glaucescens* and *M. guttatus* used in characterization of isolating barriers. All populations located in Butte County, California, USA. Experiments: 1. Flowering phenology, 2. Flower morphology, 3. Pollinator preference, 4. Pollen adhesion, 5. Pollen germination, 6. Pollen tube growth rate, 7. Intrinsic postzygotic performance. Vouchers from study populations are housed within the Chico State herbarium (CHSC).

| Species | Site name | Location | Elevation (m) | Experiments used |
|---|---|---|---|---|
| *Mimulus glaucescens* | Centerville Road | 39.73630, -121.70176 | 117 | 1, 8 |
| | Honey Run – 3 | 39.74616, -121.67212 | 173 | 1, 2, 4, 5, 6, 7 |
| | Honey Run – 4 | 39.74676, -121.65691 | 329 | 1 |
| | Honey Run – 5 | 39.74969, -121.64137 | 386 | 1, 4, 5, 6 |
| | Quail Run | 39.76781, -121.67488 | 171 | 7 |
| | Salmon Hole | 39.78321, -121.74489 | 150 | 2, 6 |
| | Sandstone Glade | 39.85254, -121.69965 | 403 | 2, 6 |
| | Simmon's Loop | 39.82230, -121.71290 | 508 | 2, 3, 6 |
| *Mimulus guttatus* | Briar Patch | 39.74162, -121.69567 | 116 | 1, 8 |
| | Bridge | 39.72743, -121.70507 | 101 | 7 |
| | Cherokee Road | 39.58420, -121.54638 | 381 | 2, 6 |
| | Cherry Hill | 40.10283, -121.50094 | 1448 | 2 |
| | Coal Canyon Road | 39.61012, -121.59636 | 161 | 2, 6 |
| | Dry Creek | 39.62168, -121.63387 | 83 | 2, 6 |
| | Honey Run – 1 | 39.74293, -121.67539 | 165 | 1, 3, 4, 5, 7 |
| | Honey Run – 2 | 39.75198, -121.63680 | 454 | 1, 4, 5, 6 |
| | Willow Springs | 39.74625, -121.68873 | 118 | 1 |

Table S2. Insects observed visiting flowers in experimental arrays of *Mimulus glaucescens* and *M. guttatus* in Butte County, California. All visitor taxa were observed on flowers of both species. Determinations by L. S. Kimsey of University of California, Davis Bohart Museum.

| Order | Family | Genus | Species |
|---|---|---|---|
| Coleoptera | Buprestidae | *Anthaxia* | sp. |
| Diptera | Anthomyiidae | | |
| | Bombyliidae | *Bombylius* | *major* |
| | Tephritidae | *Chaetorollia* | sp. |
| | Syrphidae | *Platycheirius* | *stegrus* |
| | | *Scaevis* | sp. |
| Hemiptera | Miridae | | |
| Hymenoptera | Apidae | *Apis* | *mellifera* |
| | | *Xylocopa* | *californica* |
| | Halictidae | *Lasioglossum* (*Dialictus*) | sp. |
| | Megachilidae | *Osmia* | sp. |
| Lepidoptera | Adelidae | *Adelia* | *eldorada* |

Table S5. Lower and upper 95% confidence interval boundaries, generated by 10,000 bootstrap replicates, of reproductive isolation for multiple potential barriers to reproduction between *Mimulus glaucescens* and *M. guttatus* in northern California. Estimates are provided between species as well as between populations within each species.

| Species | Barrier | Between-species | Between-populations |
|---|---|---|---|
| *M. glaucescens* | Ecogeographic* | 0.169, 0.693 | --- |
| | Flowering phenology | 0.110, 0.226 | 0.142, 0.233 |
| | Pollinator visitation | -0.296, 0.074 | --- |
| | Pollen adhesion | 0.063, 0.266 | -0.108, 0.139 |
| | Pollen germination | -0.099, 0.397 | -0.406, 0.095 |
| | Pollen-tube growth rate | -0.135, 0.192 | -0.066, 0.266 |
| | $F_1$ seed number | -0.077, 0.158 | -0.136, 0.177 |
| | $F_1$ seed germination | -0.168, 0.107 | -0.144, 0.223 |
| | $F_1$ days to first flower | 0.064, 0.369 | 0.089, 0.506 |
| | $F_1$ number of flowers | -0.071, 0.192 | 0.028, 0.363 |
| | $F_1$ ovules per carpel | -0.182, -0.062 | -0.129, 0.003 |
| | $F_1$ pollen viability | -0.230, -0.018 | -0.351, -0.165 |
| | $F_1$ plant biomass | -0.302, 0.014 | -0.039, 0.398 |
| *M. guttatus* | Ecogeographic* | 0.443, 0.716 | --- |
| | Flowering phenology | 0.148, 0.283 | 0.135, 0.256 |
| | Pollinator visitation | -0.268, 0.107 | --- |
| | Pollen adhesion | -0.226, 0.052 | -0.294, 0.129 |
| | Pollen germination | -0.046, 0.329 | -0.479, 0.006 |
| | Pollen-tube growth rate | -0.073, 0.218 | -0.068, 0.208 |
| | $F_1$ seed number | -0.070, 0.221 | -0.066, 0.208 |
| | $F_1$ seed germination | -0.154, 0.093 | -0.165, 0.116 |
| | $F_1$ days to first flower | 0.006, 0.263 | 0.003, 0.222 |
| | $F_1$ number of flowers | -0.448, -0.243 | -0.193, 0.081 |
| | $F_1$ ovules per carpel | -0.100, 0.049 | -0.087, 0.086 |
| | $F_1$ pollen viability | 0.046, 0.221 | -0.115, 0.018 |
| | $F_1$ plant biomass | -0.393, -0.139 | -0.291, 0.025 |

*1,000 bootstrap replicates were used for Ecogeographic isolation for computational efficiency

Table S4. Information on whole-genome samples used in analyses.

| Species | Sample ID | Average Coverage | Latitude | Longitude | SRA |
|---|---|---|---|---|---|
| M. caespitosa | GAB1 | 4.8 | -121.72335 | 46.88297 | SRR12424423 |
| M. caespitosa | GAB2 | 5.1 | -121.72335 | 46.88297 | SRR12424422 |
| M. caespitosa | KCK1 | 5.6 | -121.56115 | 46.84036 | SRR12424416 |
| M. caespitosa | PAG2 | 4.6 | -121.71255 | 46.79924 | SRR12424413 |
| M. caespitosa | TWN36 | 5.3 | -121.38164 | 48.57026 | SRR12424421 |
| M. caespitosa | UTC1 | 4.7 | -121.87111 | 46.8004 | SRR12424419 |
| M. caespitosa | UTC2 | 4.4 | -121.87111 | 46.8004 | SRR12424418 |
| M. corallinus | EAG | 7 | -119.92 | 38.32 | SRR13618753 |
| M. corallinus | SILC | 8.1 | -119.786 | 38.588 | SRR13618752 |
| M. corallinus | SILF | 6.6 | -120.219 | 38.664 | SRR13618751 |
| M. decorus | IMPIA | 1.7 | -122.149 | 44.3934 | SRX2211949 |
| M. decorus | IMPO | 19.5 | -122.149 | 44.3934 | SRR10194630 |
| M. decorus | (DL1) S1 | 8.5 | -122.133433 | 43.1575333 | SRR10194633 |
| M. decorus | HACK (S10) | 5.6 | -122.033833 | 45.60705 | SRR10194634 |
| M. decorus | LL (S2) | 8.6 | -122.0542 | 44.1739333 | SRR10194642 |
| M. decorus | HWY15 (S3) | 7.8 | -122.1 | 44.2 | SRR10194631 |
| M. decorus | BR5 (S5) | 7.1 | -122.104 | 44.371 | SRR10194636 |
| M. decorus | Odell-11 (S6) | 8.6 | -121.962783 | 43.54795 | SRR10194639 |
| M. decorus | HJA2 (S7) | 7.5 | -122.1762 | 44.2332 | SRR10194632 |
| M. decorus | KINK (S80 | 6.4 | -121.996217 | 44.3032 | SRR10194643 |
| M. glaucescens | HRH16 | 4.6 | -121.64137 | 39.74969 | TBA |
| M. glaucescens | HRL21 | 6 | -121.67212 | 39.74616 | TBA |
| M. glaucescens | DC | 2.2 | -121.712967 | 39.8223 | TBA |
| M. glaucescens | PMO | 6 | 121.6336 | 39.96713333 | TBA |
| M. glaucescens | PUL5 | 6.6 | -121.452817 | 39.79523 | TBA |

| | | | | | |
|---|---|---|---|---|---|
| M. glaucescens | RED | 1.5 | -121.704217 | 39.72746667 | TBA |
| M. laciniatus | OPN | 6.7 | | | TBA |
| M. laciniatus | Per | 4.9 | -119.3687 | 37.055767 | TBA |
| M. laciniatus | TRT | 5.8 | -119.70535 | 37.7165 | TBA |
| M. laciniatus | WLF | 2.4 | -119.59385 | 37.841533 | SRR071764 |
| M. minor | NOR511 | 6.2 | -107.045033 | 39.015033 | SRR12424415 |
| M. minor | NOR523 | 5.7 | -107.045033 | 39.015033 | SRR12424414 |
| M. minor | UNP12 | 5.3 | -107.096053 | 39.019639 | SRR12424420 |
| M. nasutus | CACN9 | 10.1 | -121.3667 | 45.71076 | SRR1259271 |
| M. nasutus | DPRN104 | 9.5 | -120.344 | 37.828 | SRR1259273 |
| M. nasutus | KOOT | 12.7 | -115.983 | 48.104 | SRR1259272 |
| M. nasutus | NHN26 | 9.8 | -124.16 | 49.273 | SRX525051 |
| M. nasutus | SF5 | 3.4 | -121.0225 | 45.264444 | SRX116529 |
| M. pardalis | DPP2 | 9.2 | -120.41745 | 37.700059 | SRR1298375 |
| M. pardalis | SEC39 | 4.1 | -118.5767 | 42.6364 | SRR9674910 |
| M. tilingii | A25 | 1.9 | -118.5767 | 42.6364 | SRX6914883 |
| M. tilingii | LVR | 13.9 | -119.13544 | 37.57049 | SRX1532174 |
| M. tilingii | ICE10 | 4.7 | -117.16067 | 45.13553 | SRR12424417 |
| M. tilingii | SAB1 | 5.2 | -118.36627 | 37.1272 | SRR12424411 |
| M. tilingii | SAB19 | 4.6 | -118.36627 | 37.1272 | SRR12424412 |
| M. tilingii | SOP12 | 5.2 | -119.617981 | 38.266753 | SRR12424410 |
| Northern M. guttatus | BP9 | 5.3 | -121.69567 | 39.74162 | TBA |
| Northern M. guttatus | HRH10 | 5.6 | -121.6368 | 39.75198 | TBA |
| Northern M. guttatus | AHQT | 7 | -110.813 | 44.431 | SRX142379 |
| Northern M. guttatus | ALK4 | 1.5 | -157.36 | 57.2 | SRX6914884 |
| Northern M. guttatus | BOG | 3.9 | -118.805833 | 41.923611 | SRX030570 |
| Northern M. guttatus | CACG6 | 13.9 | -121.3667 | 45.71076 | SRX525044 |
| Northern M. guttatus | CG | 1.9 | | | SRX6914893 |
| Northern M. guttatus | IM767 | 3.3 | -122.508783 | 45.57571 | SRX487581 |

| | | | | | |
|---|---|---|---|---|---|
| Northern M. guttatus | MAR | 3.8 | -123.29445 | 43.4786 | SRX030542 |
| Northern M. guttatus | SOL | 2.6 | -119.175168 | 41.378995 | SRX6914890 |
| Northern M. guttatus | TSG3 | 1.7 | -131.91573 | 53.418833 | SRX2019854 |
| Northern M. guttatus | YJS6 | 3.1 | -114.5845 | 44.9512 | SRX030545 |
| Northern M. guttatus | ATTU | 4.9 | 55.46 | -4.63 | SRX10011990 |
| Northern M. guttatus | GUT5 | 4.4 | -118.57 | 42.71 | SRX10011991 |
| Southern M. guttatus | Odell_gutt(S11) | 8 | -121.962783 | 43.54795 | SRR10194640 |
| Southern M. guttatus | CSS4 | 4.9 | -122.415278 | 38.861111 | SRX6435296 |
| Southern M. guttatus | INV | 3.8 | -122.87 | 38.08 | SRX6914899 |
| Southern M. guttatus | LMC24 | 3 | -123.083917 | 38.863983 | SRX030680 |
| Southern M. guttatus | MED84 | 9.7 | -120.313667 | 37.816633 | SRX552649 |
| Southern M. guttatus | MEX | 2.2 | -111.09455 | 29.101357 | SRX6914887 |
| Southern M. guttatus | REM8 | 2.8 | -122.411467 | 38.860433 | SRX030546 |
| Southern M. guttatus | SCH | 10.1 | -107.035267 | 39.018467 | SRX371892 |
| Southern M. guttatus | SLP | 8.7 | -120.461869 | 37.84825642 | SRX142377 |
| Southern M. guttatus | SWB | 3.7 | -123.690467 | 39.035983 | SRX030679 |
| Southern M. guttatus | YV06 | 6.5 | -119.746433 | 37.723367 | SRX6914891 |
| Southern M. guttatus | DUN | 1.6 | 43.893 | -124.138 | SRR072711, SRR072713 |
| Southern M. guttatus | NRM | 3.2 | -105.675589 | 40.334795 | SRX10011992 |
| Southern M. guttatus | PED5 | 3.2 | -110.78417 | 32.982928 | SRR071969 |
| Southern M. guttatus | SHG | 5.1 | -122.415278 | 38.861111 | SRX10011994 |

Table S5. Pairwise estimates of $F_{ST}$ (above the diagonal) and $D_{XY}$ (below the diagonal) across *Mimulus* taxa sampled, based on analyses of whole-genome

| $F_{ST}$/$D_{XY}$ | M. glaucescens | Southern M. decorus | Northern M. decorus | Northern M. guttatus | M. corallinus | Southern M. guttatus | M. pardalis | M. laciniatus | M. nasutus | M. tilingii | M. caespitosa |
|---|---|---|---|---|---|---|---|---|---|---|---|
| M. glaucescens | 0.0328551 | 0.13782269 | 0.38756289 | 0.08015987 | 0.18412838 | 0.11829728 | 0.23075542 | 0.3805836 | 0.47605777 | 0.36800694 | 0.5746358[illegible] |
| Southern M. decorus | 0.04394185 | 0.0380263 | 0.22292097 | 0.05791609 | 0.17325247 | 0.08969018 | 0.25621103 | 0.44328939 | 0.57089703 | 0.35347488 | 0.645612[illegible] |
| Northern M. decorus | 0.04182793 | 0.03115866 | 0.01816435 | 0.31145644 | 0.44667097 | 0.32809987 | 0.5615156 | 0.62164449 | 0.67371337 | 0.55078675 | 0.6998833[illegible] |
| Northern M. guttatus | 0.03927803 | 0.04141665 | 0.04036643 | 0.03541647 | 0.14644489 | 0.10402327 | 0.20268826 | 0.33290011 | 0.39241923 | 0.33209729 | 0.4772952[illegible] |
| M. corallinus | 0.04447217 | 0.04433555 | 0.04450847 | 0.04506125 | 0.03788315 | 0.08000938 | 0.23083959 | 0.38947009 | 0.50220404 | 0.34288324 | 0.6012559[illegible] |
| Southern M. guttatus | 0.04337536 | 0.04585161 | 0.04410567 | 0.04423352 | 0.04268013 | 0.03854437 | 0.10915211 | 0.23254555 | 0.29807115 | 0.30214061 | 0.4488888[illegible] |
| M. pardalis | 0.04330223 | 0.04631394 | 0.04498966 | 0.044201 | 0.04201778 | 0.04036189 | 0.02104308 | 0.46435658 | 0.61508523 | 0.44673032 | 0.7430538[illegible] |
| M. laciniatus | 0.04520601 | 0.04741692 | 0.04646721 | 0.04532664 | 0.04382875 | 0.04180177 | 0.04079296 | 0.01597 | 0.52489291 | 0.5491206 | 0.7559122[illegible] |
| M. nasutus | 0.04609177 | 0.04649176 | 0.04682451 | 0.04331372 | 0.04616263 | 0.04243891 | 0.04260426 | 0.03116571 | 0.00921865 | 0.62526226 | 0.8012529[illegible] |
| M. tilingii | 0.04588754 | 0.04508422 | 0.0431607 | 0.04732023 | 0.04436008 | 0.04572103 | 0.04503947 | 0.04586247 | 0.04745273 | 0.02041857 | 0.5601138 |
| M. caespitosa | 0.04565267 | 0.04473115 | 0.04223606 | 0.04508915 | 0.04615279 | 0.04680066 | 0.046758 | 0.04663902 | 0.04713941 | 0.03480894 | 0.0079820[illegible] |
| M. minor | 0.04770525 | 0.04653758 | 0.04443498 | 0.04908843 | 0.04741432 | 0.04786507 | 0.04767135 | 0.04881907 | 0.05108073 | 0.03518768 | 0.031377[illegible] |

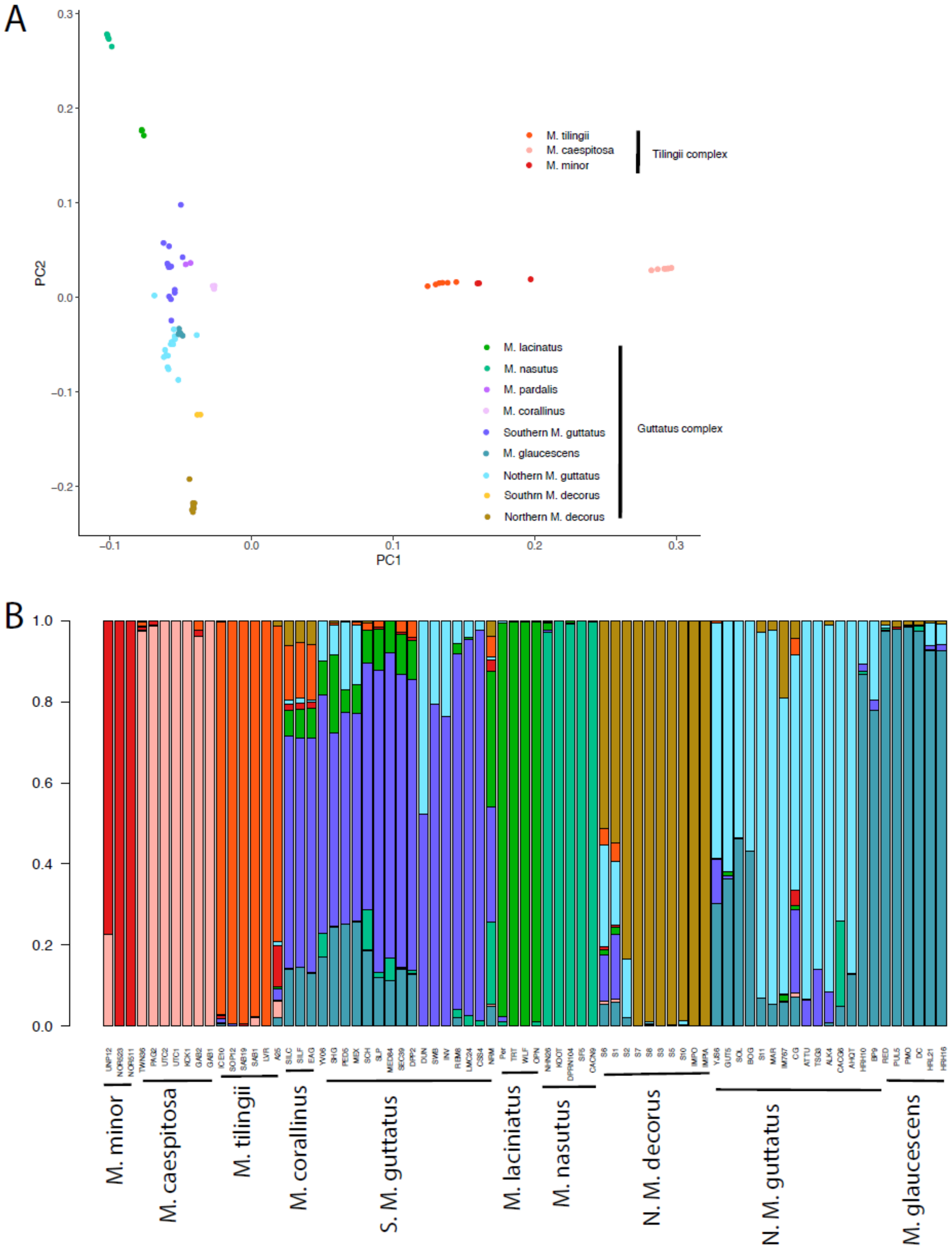
A
PC1
PC2
M. tilingii
M. caespitosa
M. minor
Tilingii complex
M. lacinatus
M. nasutus
M. pardalis
M. corallinus
Southern M. guttatus
M. glaucescens
Nothern M. guttatus
Southrn M. decorus
Northern M. decorus
Guttatus complex
B
M. minor
M. caespitosa
M. tilingii
M. corallinus
S. M. guttatus
M. laciniatus
M. nasutus
N. M. decorus
N. M. guttatus
M. glaucescens

Figure S1. (A) First two principal components of whole-genomic variation in *Mimulus* accessions used in study. (B) Proportion of genome associated with ancestry of *Mimulus* taxa sampled in study (see also Table S3) based on K-means clustering analysis. Colors correspond to taxa indicated in A.

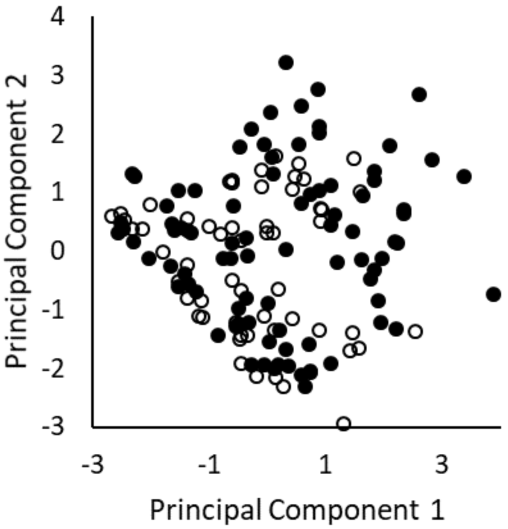


Figure S2. Scores of the first two principal components of six individual-plant scale habitat characteristics of *Mimulus glaucescens* (filled circles) and *M. guttatus* (open circles) growing in a 6.0 $km^2$ region of sympatry within Butte County, California, USA.

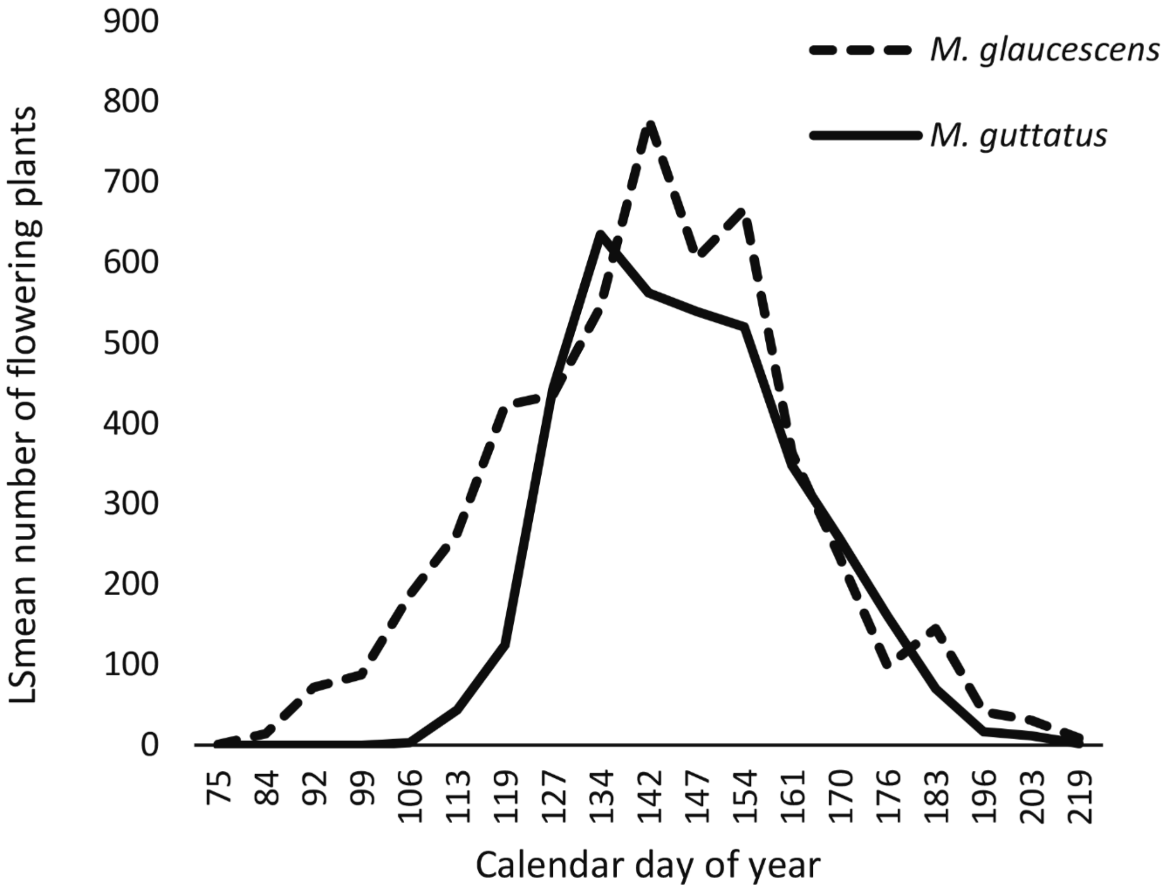


Figure S3. Least-squares mean number of flowering plants from weekly census counts of four sympatric populations each of *Mimulus glaucescens* and *M. guttatus* occurring in Butte County, California, USA.

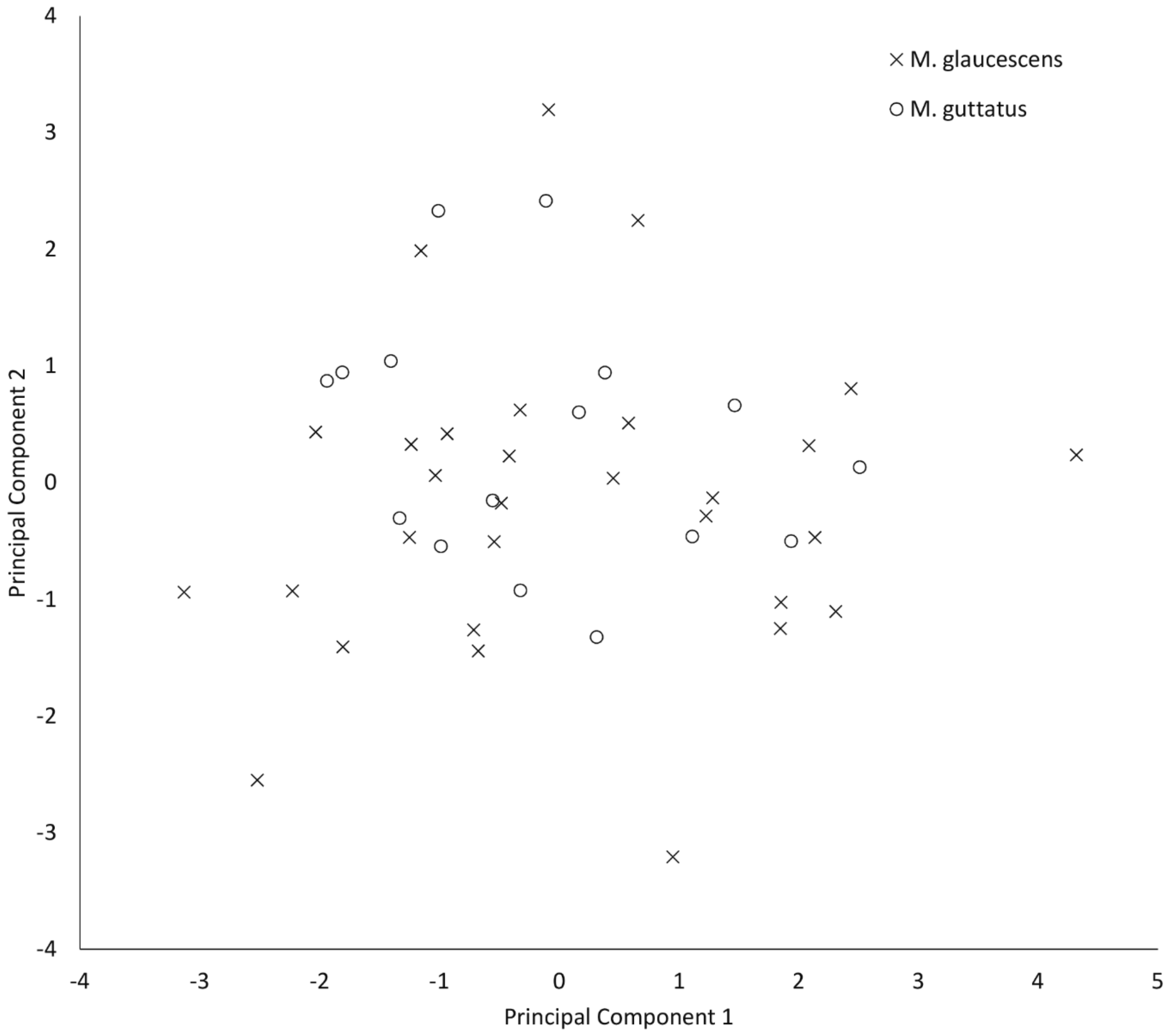


Figure S4. Scores of the first two principal components based on family mean values of floral morphological traits for *Mimulus glaucescens* and *M. guttatus* grown in the greenhouse from open-pollinated seeds collected in populations throughout Butte County, California, USA.

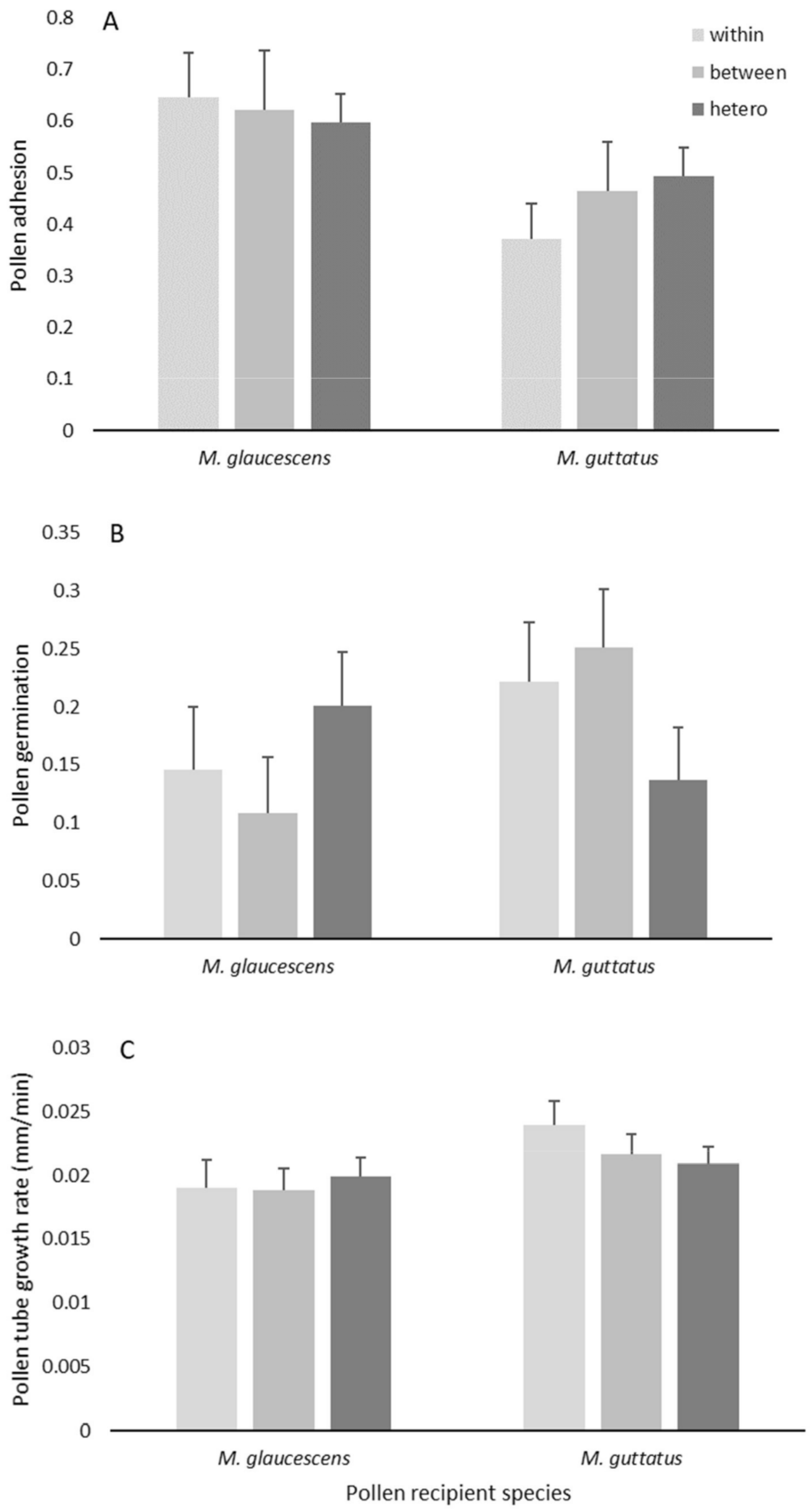

A
within
between
hetero
Pollen adhesion
0.8
0.7
0.6
0.5
0.4
0.3
0.2
0.1
0
M. glaucescens
M. guttatus
B
Pollen germination
0.35
0.3
0.25
0.2
0.15
0.1
0.05
0
M. glaucescens
M. guttatus
C
Pollen tube growth rate (mm/min)
0.03
0.025
0.02
0.015
0.01
0.005
0
M. glaucescens
M. guttatus
Pollen recipient species

Figure S5. Least-squares mean proportion of pollen grains adhering to stigmas (A), proportion of adhered pollen grains successfully germinating (B), and pollen-tube growth rate within styles of experimental hand-pollinations of *Mimulus glaucescens* and *M. guttatus* conducted between plants derived from the same population ("within"), between plants of the same species but from different populations ("between"), and between species ("hetero"). Error bars represent 1 S.E. All plants were grown in the greenhouse from field-collected open-pollinated seeds. See Methods for additional information about experimental protocols.

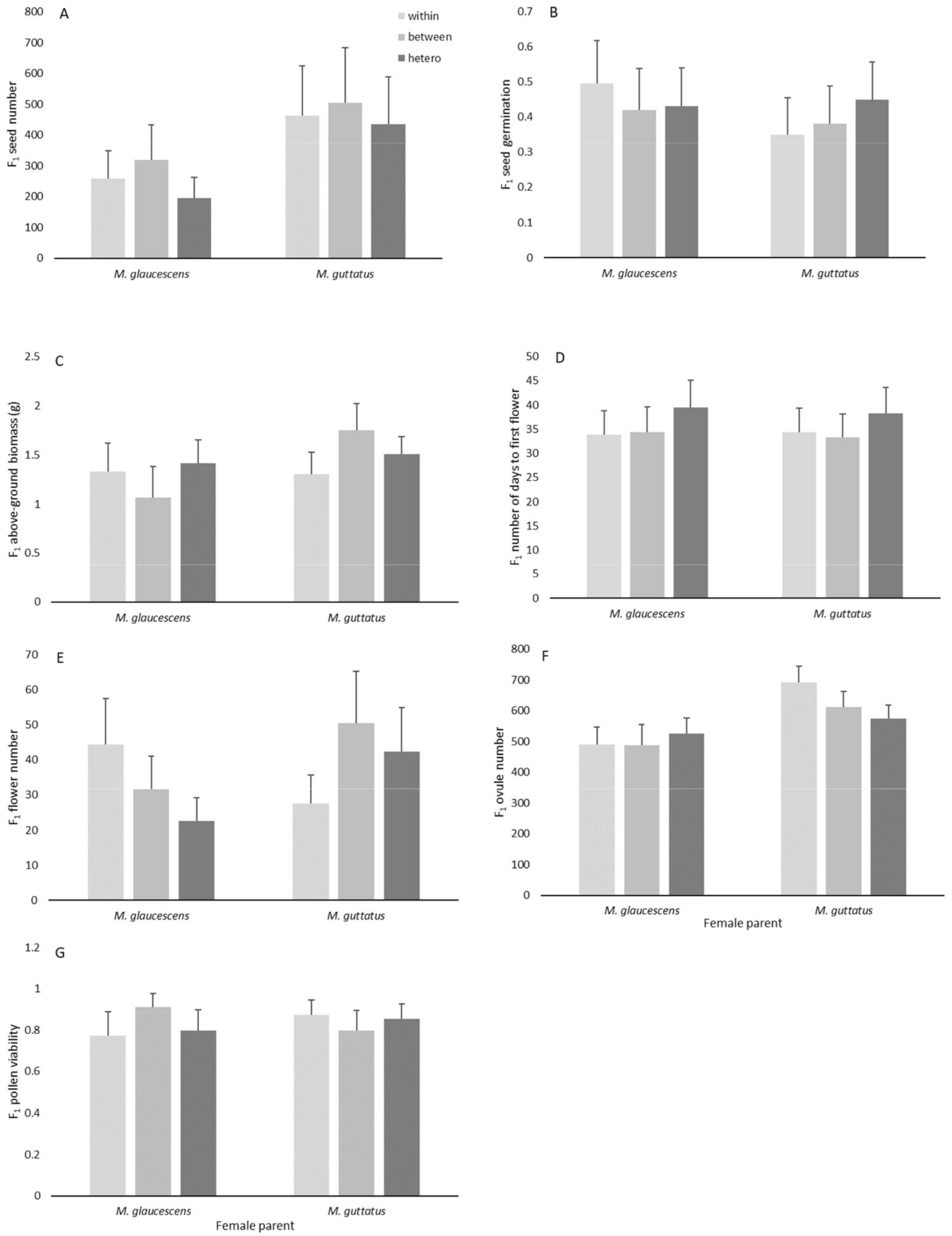


Figure S6. Performance of $F_1$ offspring produced from crosses between *Mimulus glaucescens* and *M. guttatus* originating from open-pollinated seeds collected in sympatric populations. Error bars represent 1 S.E. Parents and $F_1$s were reared in laboratory cultivation (see methods). Crosses included plants derived from the same population (“within”), between plants of the same species but from different populations (“between”), and between species (“hetero”). Least-squares mean (A) number of seeds per fruit derived from $F_1$ crosses, (B) proportion of seeds germinating, (C) above-ground dry biomass, (D) number of days to first flower anthesis, (E) number of flowers produced, (F) number of ovules per carpel, and (G) proportion of viable pollen grains.
